# Developmental single-cell transcriptomics of hypothalamic POMC progenitors reveal the genetic trajectories of multiple neuropeptidergic phenotypes

**DOI:** 10.1101/2021.05.12.443898

**Authors:** Hui Yu, Marcelo Rubinstein, Malcolm J Low

## Abstract

Proopiomelanocortin (POMC) neurons of the hypothalamic arcuate nucleus are essential to regulate food intake and energy balance. However, the ontogenetic transcriptional programs that specify the identity and functioning of these neurons are poorly understood. Here, we use scRNAseq to define the transcriptomes characterizing POMC progenitors in the developing hypothalamus and TRAP-seq to analyze the subsequent translatomes of mature POMC neurons. Our data show that *Pomc*-expressing neurons originate from multiple developmental pathways expressing distinct combinations of transcription factors. The predominant cluster, featured by high levels of *Pomc* and *Prdm12* transcripts represents the precursor of canonical arcuate POMC neurons. Additional clusters of progenitors expressing lower levels of *Pomc* mature into different neuronal phenotypes characterized by distinct combinations of transcription factors, neuropeptides, processing enzymes, cell surface and nuclear receptors. We conclude that the genetic programs specifying the identity and differentiation of arcuate POMC neurons are diverse and generate a heterogeneous repertoire of neuronal phenotypes early in development that continue to evolve postnatally.

## Introduction

*Proopiomelanocortin (POMC)*-expressing neurons located in the arcuate nucleus of the hypothalamus play an essential role in the regulation of food intake by maintaining a melanocortin-dependent anorexigenic tone. The critical importance of central melanocortins, specifically in arcuate neurons, is apparent in mice lacking *Pomc* that develop hyperphagia and early-onset extreme obesity(Bumaschny et al., 2012). This phenotype is closely mirrored in humans carrying biallelic null-mutations in *POMC*(Krude et al., 1998). In addition, arcuate POMC neurons release the *Pomc*-encoded opioid peptide β-endorphin, which is critical in mediating normal food intake and stress-induced analgesia(Parikh et al., 2011).

Postmitotic POMC precursors acquire their phenotypic identity in the developing hypothalamus days earlier than any other peptidergic neurons born in the presumptive arcuate nucleus (eg. NPY/AGRP, GHRH, SST, KISSPEPTIN). *Pomc* mRNA is initially observed in the tuberal portion of the prospective mouse hypothalamus of E10.5 embryos, following the concurrent advent of the essential transcription factors (TFs) ISL1(Nasif et al., 2015), NKX2-1(Orquera et al., 2019) and PRDM12(Hael, Rojo, Orquera, Low, & Rubinstein, 2020). In fact, the early ablation of either *Isl1*, *Nkx2-1* or *Prdm12* disrupts the onset of hypothalamic *Pomc* expression in conditional mutant mouse embryos(Hael et al., 2020; Nasif et al., 2015; Orquera et al., 2019). Given that the number of arcuate neurons coexpressing *Nkx2-1*, *Isl1* and *Prdm12* greatly exceeds that of POMC neurons in this area, it is expected that additional and still unknown TFs integrate the complete genetic program that specifies the early identity of POMC neurons.

Other molecular markers of arcuate POMC neurons such as transporters, receptors, channels and co-transmitters exhibit great variability indicating that a heterogeneous pool of diverse subpopulations of these neurons are involved in multiple circuits and functions. In fact, previous studies have suggested that subsets of arcuate *Pomc-*expressing neurons may act as precursors of terminally differentiated neurons of alternative neuropeptidergic phenotypes such as NPY/AGRP and KISSPEPTIN/NEUROKININ-B/DYNORPHIN (KNDY) neurons in the adult hypothalamus(Padilla, Carmody, & Zeltser, 2010; Sanz et al., 2015). Initial attempts to characterize the transcriptome of adult POMC neurons at the single-cell level confirmed the heterogeneous nature of arcuate POMC neurons and revealed the existence of distinct clusters including those co-expressing the aforementioned neuropeptide genes(Campbell et al., 2017; R. Chen, Wu, Jiang, & Zhang, 2017; Huisman et al., 2020; Lam et al., 2017). However, the molecular signatures of the precursors of the various hypothalamic *Pomc*-expressing lineages and their transition into mature neurons within the postnatal Arc are still lacking.

Here, we track the origin and maturation of hypothalamic POMC precursors in the developing and early postnatal hypothalamus by performing single-cell RNA-seq transcriptomics of fluorescently labeled cells taken from *Pomc-TdDiscomaRed-Sv40PolyA* (*Pomc-TdDsRed*) transgenic mice at different embryonic (E11.5, E13.5, E15.5 and E17.5) and early postnatal (P5 and P12) ages. Our data suggested that the POMC precursors arise from multiple different transcriptional routes. Some of these precursors further develop into mature POMC neurons while others follow diverse neuropeptidergic trajectories indicating that a heterogeneous population of *Pomc*-expressing neurons is established early in the developing arcuate nucleus. Combining translating ribosome affinity purification with RNA-sequencing (TRAP-seq) of *Pomc-eGFPL10a* transgenic mice at P12 and P60, we defined transcriptional programs that function distinctly in early embryonic POMC precursors and later in mature POMC neurons. These datasets provide valuable resources for comprehensively understanding the genetic development of POMC neurons.

## Results

### Cluster analysis of gene expression in Pomc positive hypothalamic cells integrated across six development ages

We captured the single cell transcriptomes of hypothalamic *Pomc*-expressing cells at four key embryonic days (E11.5, E13.5, E15.5 and E17.5) and two early postnatal days (P5 and P12) when the maturing neurons already develop axons and innervate key target nuclei distal to the Arc(Bouret, Bates, Chen, Myers, & Simerly, 2012; Bouret, Draper, & Simerly, 2004a, 2004b). Medial basal hypothalami from mice that express the reporter transgene *Pomc-TdDsRed* selectively in POMC cells were pooled from multiple embryos or postnatal pups at the indicated ages. Dissociated single cell suspensions were sorted for DsRed fluorescence and used for library preparation with the 10X genomics platform (Figure 1A). After a rigorous screening procedure to remove substandard scRNA sequencing data (see Methods), we used a combination of computational platforms to analyze the transcriptomes of 13,962 cells that had *Pomc* counts of ≥ 1 unique molecular identifiers (UMI).

**Figure 1.**
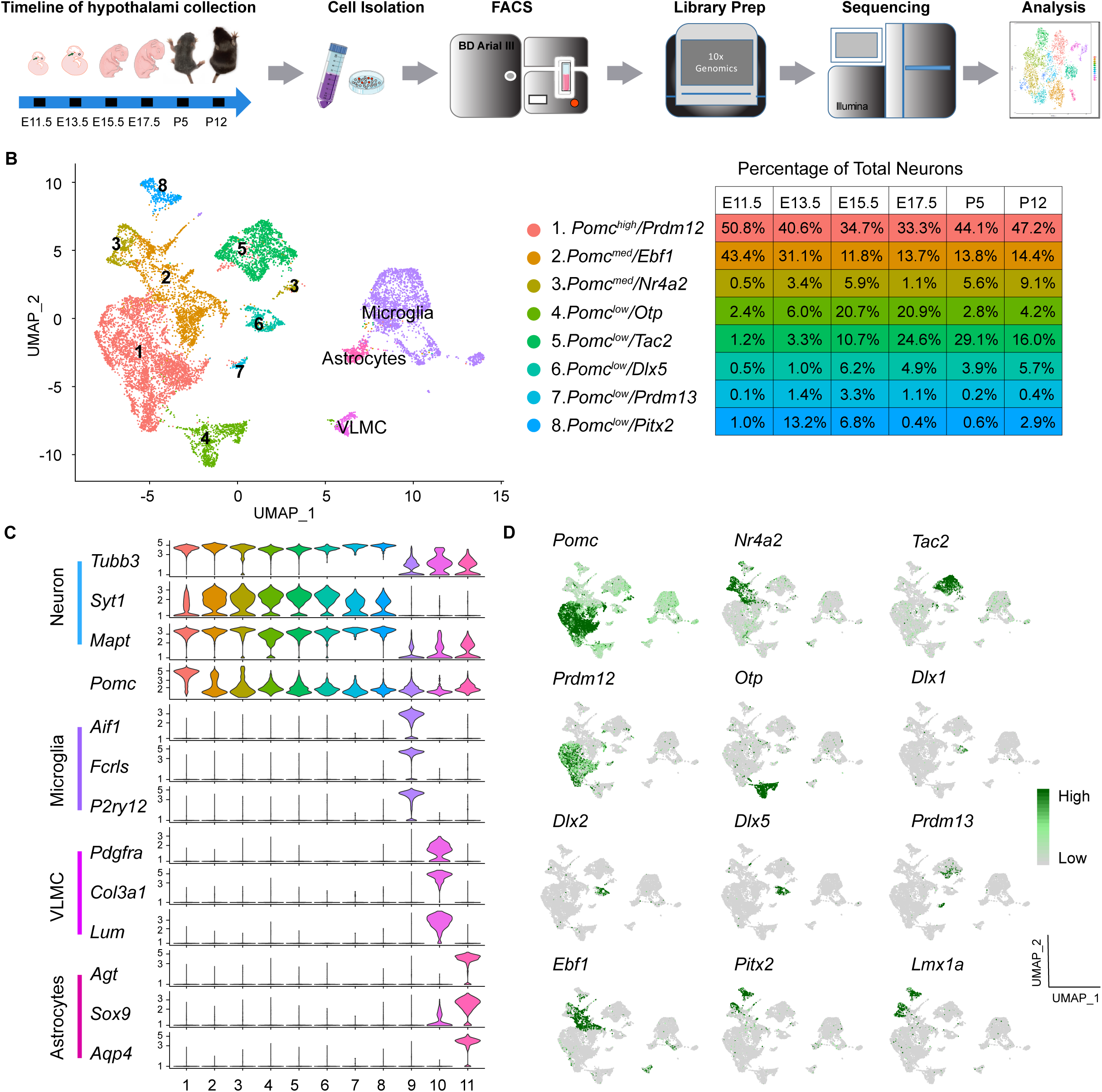
Single cell sequencing of *Pomc^DsRed^* cells from developing mouse hypothalami at E11.5, E13.5, E15.5, E17.5, P5 and P12 reveals eight distinct neuronal clusters and three non-neuronal clusters. (A) Schematic diagram showing the overall experimental procedure. (B) UMAP (Uniform Manifold Approximation and Projection) plot showing the distribution of major cell clusters and their corresponding cell% at each developmental stage. (C) Violin plots showing the expression of three signature genes and *Pomc* for each cluster. (D) UMAP plots showing the enrichment of feature genes in each cluster. Genes representing each cluster were colored and highlighted to show the cluster-specific enrichment. The color intensity corresponds to the normalized gene expression from Seurat analysis, where gene expression for each cell was normalized by the total expression, multiplied by a scale factor of 10,000, and then log-transformed. VLMC, vascular and leptomeningeal cell.

A Seurat analysis of all cells integrated across the six-time points revealed 11 distinct cell clusters projected onto a UMAP plot (Figure 1B). The same data are represented in a heat map corresponding to the top 30 expressed genes per cluster (Supplementary Figure 1). Clusters 1-8 (10,828 cells) were all highly enriched for a panel of three neuronal marker gene transcripts. The embedded table in Figure 1B lists the percentages of neurons of each cluster at each developmental age. The other three clusters (3,134 cells) were enriched in transcripts characteristic of either microglia, vascular and leptomeningeal cells (VLMC) or astrocytes (Figure 1C). GO ontology analyses for biological processes further confirmed the neuronal or non-neuronal identities of the 11 individual clusters (data not shown).

By definition, all cells in the neuronal and non-neuronal clusters contained *Pomc* transcripts (UMI ≥ 1), however there were significant differences in the average expression levels per cluster, ranging from 1.3 to 33.8 [exp(*log Normalized* counts) -1)]. Nomenclature of the 8 neuronal clusters is based on a combination of relatively high (>20), medium (5-20) or low (<5) *Pomc* levels and high expression of one differential key feature gene as follows: no. 1, *Pomc^high^/Prdm12*; no. 2 *Pomc^med^/Ebf1*; no. 3, *Pomc^med^/Nr4a2*; no. 4, *Pomc^low^/Otp*; no. 5, *Pomc^low/^Tac2*; no. 6, *Pomc^low^/Dlx5*; no. 7, *Pomc^low^/Prdm13*; and no. 8, *Pomc^low^/Pitx2*. The cluster distribution and relative expression of those feature genes are represented by individual UMAP plots (Figure 1D). Average expression levels of all genes, including *Pomc*, in the 11 clusters and complete lists of all feature genes that define the individual clusters are presented in Supplemental Tables 1 and 2, respectively.

In order to validate our experimental approach of isolating *Pomc*-expressing cells from the developing hypothalamus by FACS for the transgenic surrogate marker *Pomc-TdDsRed*, we performed an independent unsupervised cluster analysis of the transcriptomes from the 5,724 cells that had *DsRed* counts of ≥ 1 UMI (Supplemental Figure 2A-C and Supplemental Table 3). A projection analysis (Supplemental Figure 2D-F) of the differentially expressed feature genes defined 9 DsRed clusters (Supplemental Table 4) which closely corresponded to 9 of the 11 clusters based on *Pomc* counts of ≥ 1 UMI (Supplemental Table 2). The two exceptions were the smallest clusters 7. *Pomc^low^/Prdm13* and Astrocytes (Supplemental Figure 2D-F). Furthermore, the percentages of cells at each developmental age from the corresponding neuronal clusters were remarkably similar in the *Pomc* UMI ≥ 1 data set compared to the *DsRed* UMI ≥ 1 data set (compare Figure 1B to Supplemental Figure 3A). The different population of DsRed fluorescent sorted cells showing UMI ≥ 1 for *Pomc* (13,962) or *DsRed* (5,724) may be due to differential half-lives of the fluorescent protein DSRED used during cell sorting and its transcript used for barcoding the RNAseq libraries.

### Temporal gene expression analysis of the eight neuronal clusters at individual developmental stages

Next, we analyzed temporal gene expression patterns for each of the 8 neuronal clusters at the 6 distinct developmental ages (Supplemental Table 5). The progressive changes in feature genes across time are responsible for the observed within cluster heterogeneity on the heat map in Supplemental Figure 1.

#### Cluster 1

*Pomc^high^/Prdm12* cells were the most abundant among the eight neuronal clusters and consistently had the highest levels of *Pomc* expression at all six developmental ages. We consider them to be the direct precursors of canonical POMC neurons in the adult hypothalamus (Figure 2A-2C). Other feature genes included a set of four TFs known to be critical for the initiation and/or maintenance of Arc *Pomc* expression, *Isl1, Nkx2-1, Prdm12* and *Tbx3(Hael et al., 2020; Nasif et al., 2015; Orquera et al., 2019; Quarta et al., 2019)* (Figure 2A). They were expressed in similar developmental patterns to each other, with the exception of *Tbx3* that was significantly expressed only from day E15.5. Notably, there was a lack of the pituitary-specific *Tbx19* TF, confirming that no pituitary tissue was included in the hypothalamic dissections. Two homologues of the *Drosophila sine oculus* homeobox gene, *Six3* and *Six6,* which have not been described previously in the context of Arc POMC neuron development, were also expressed robustly at all six time points (Figure 2A). Co-expression of their mRNAs with *Pomc* by RNAScope at ages E12.5 and E16.5 confirmed the scRNAseq analysis, although the two TFs were expressed in additional hypothalamic domains and the pituitary gland (Figure 2C). *Vim1, Sox1, 2, 3 and 14,* and *Myc*, whose expression is typically associated with neuronal precursors or immature neurons, tended to have higher expression at the earlier embryonic ages (Figure 2A). The nuclear receptor gene *Nr5a1*, also known as steroidogenic factor 1 (*Sf1*), was a key feature gene expressed almost exclusively at ages E11.5. (Figure 2A), but only minimally expressed at later embryonic time points when NR5A1 is critically involved in the development of the hypothalamic ventromedial nucleus (VMN)(Ikeda, Luo, Abbud, Nilson, & Parker, 1995). This differential spatio-temporal presence of NR5A1 within or outside POMC neurons was confirmed by immunofluorescence on hypothalamic sections taken from E11.5 and E17.5 mouse embryos (Figure 2C). Interestingly, *Nr5a1*, *Nr0b1* (*Dax1*) and *Shh*, transcripts were also present in Cluster 1 at the two earliest ages, and are factors known to interact with the *Wnt/*β -catenin signaling pathway in early cell fate determination in the adrenal and pituitary glands(Luo, Ikeda, & Parker, 1994; Meeks, Weiss, & Jameson, 2003; Mizusaki et al., 2003). A combination of pseudotime and gene ontology analyses for the top expressed genes (Log2FoldChange >0.25, *P*<0.05) in cluster1 revealed that the genes expressed at the beginning of pseudotime function in the organization of cell projections, axonogenesis and the regulation cellular component size, whereas genes expressed at the middle and end of the pseudotime function are related to neuron maturation and neuron-specified functions such as the formation of myelin sheath and regulation of synaptic vesicle cycling (Figure 2B).

**Figure 2.**
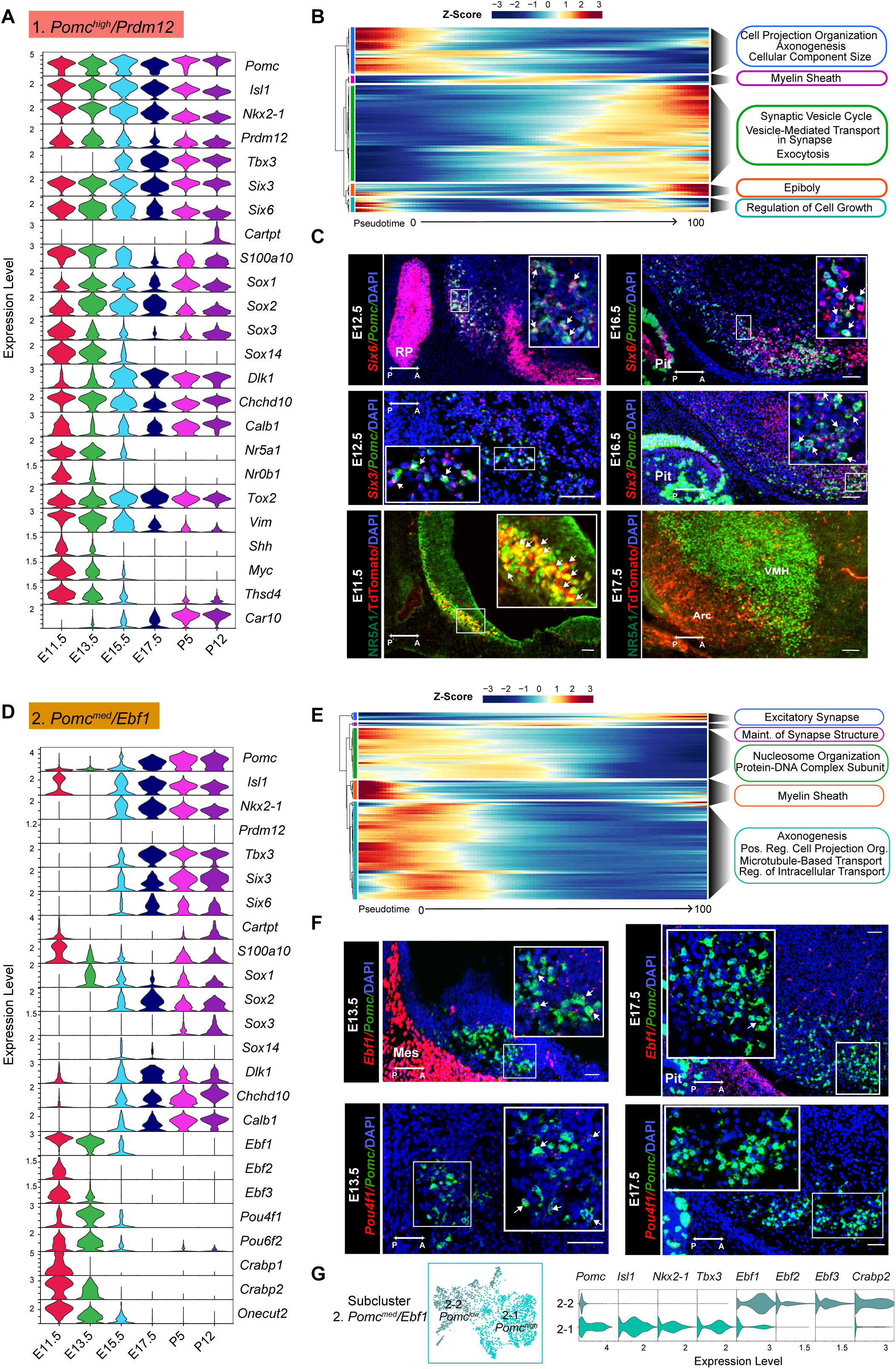
Temporal gene expression patterns of the *Pomc^high^/Prdm12* cluster and *Pomc^med^/Ebf1* clusters. (A) and (D) Violin plots showing the expression of signature genes at each developmental stage in the *Pomc^high^/Prdm12* cluster and *Pomc^med^/Ebf1* cluster respectively. (B) and (E) Heatmaps showing the gene ontology analysis based on the top marker genes in the order of pseudotime in *Pomc^high^/Prdm12* cluster and *Pomc^med^/Ebf1* cluster respectively. (C) Fluorescence *in situ* hybridization showing the co-localization of *Pomc* (green) and *Six6* (red), the co-localization of *Pomc* (green) and *Six3* (red) at indicated developmental stages. Immunofluorescence showing the co-localization TDTOMATO (red) NR5A1 (green) at the indicated developmental stages. (F) Fluorescence *in situ* hybridization showing the co-localization of *Pomc* (green) and *Ebf1* (red) and the co-localization of *Pomc* (green) and *Pou4f1* (red) at indicated developmental stages. (G) Re-clustering cells from the *Pomc^med^/Ebf1* cluster reveals two sub-clusters denoted as 2-1 *Pomc^high^* and 2-2 *Pomc^low^*. Violin plots showing the signature gene expression in 2-1 *Pomc^high^* and 2-2 *Pomc^low^* clusters. Insets are magnified views of the indicated boxes. Arrows indicate co-expressing neurons in the merged panels. Scale bar: 50 µm. Image orientation: left, posterior(P); right, anterior(A). RP: Rathke’s pouch; Pit: pituitary gland. Arc: arcuate nucleus; VMH: ventromedial hypothalamus. Mes: mesenchyme.

#### Cluster 2

*Pomc^med^/Ebf1* neurons made up the second highest abundance cluster of POMC neurons, particularly at E11.5 and E13.5. After these time points, the fraction of all neurons constituting cluster 2 dropped by approximately two thirds from ages E15.5 through P12. In contrast, the relative percentage of neurons in clusters 3-8 increased over time with peaks at different embryonic ages (Figure 1B). Cluster 2 neurons, on average, had medium levels of *Pomc* transcripts that were divided distinctly from their lowest at ages E11.5-E13.5 to highest at later embryonic and postnatal ages (Figure 2D; Supplemental Table 5). Major cluster 2 feature markers included members of the non-basic helix-loop-helix early B-cell TF family EBF (Supplemental Table 2). *Ebf1* is typically co-expressed with either *Ebf2* or *Ebf3* and pairs of these factors were shared not only in cluster 2, but also in clusters 3 and 8 (Supplemental Figure 4A and D). Dual RNAScope *in situ* hybridization showed overlap of *Ebf1* and *Pomc* mRNAs in hypothalamic neurons at an early embryonic stage (E13.5), whereas at later embryonic stage (E17.5), POMC neurons had barely detectable *Ebf1* mRNAs (Figure 2F). A similar expression pattern was found with another cluster 2 featured gene, *Pou4f1 (POU Class 4 Homeobox 1)*. Dual RNAScope *in situ* hybridization confirmed the overlap of *Pou4f1* and *Pomc* mRNAs at E13.5, but not at E17.5 (Figure 2F). These results, together, suggest that *Pomc* mRNA levels peak only after expression of both *Ebf1* and *Pou4f1* decrease. Two other prominent feature genes in cluster 2 were *Crabp1* and *Crabp2*, which encode cellular retinoic acid binding proteins. A majority of the marker genes for cluster 2 are predicted to be expressed at the beginning of pseudotime and the main functions are associated with the maintenance of basic cell biological processes such as nucleosome organization, axonogenesis, microtubule transport, and intracellular transport (Figure 2E). The genes ordered in the middle and end of pseudotime function for the regulation of synapse structure and activities.

Visual examination of the UMAP plots in Figure 1D suggested that cluster 2 was composed of two compartments, one with high and one with low *Pomc* abundance. Unsupervised reclustering of the differentially expressed genes from integrated cluster 2 confirmed the existence of two subclusters (Figure 2G). Subcluster 2-1 contained two-thirds of the neurons, which were *POMC^high^* and present only from ages E15.5-P12. Subcluster 2-2 contained the other one-third of cluster 2 neurons, which in contrast were *Pomc^low^* and detected at ages E11.5 and E13.5. Consistently, known activating TFs for *Pomc* expression such as *Nkx2-1*, *Isl1* and *Tbx3* were limited to subcluster 2-1 while the *Ebf* and *Crabp* transcripts were characteristic of subcluster 2-2 (Supplemental Table 6 and Supplemental Table 7). Conspicuously absent from both subclusters was *Prdm12* mRNA.

#### Cluster 3

*Pomc^med^/Nr4a2* neurons displayed high expression of its major feature gene *Nr4a2* (Nuclear Receptor Subfamily 4 Group A Member 2) (Supplemental Figure 4A). *Nr4a2* mRNA was also expressed at relatively high levels in clusters 2. *Pomc^med^/Ebf1* and 8. *Pomc^low^/Pitx2*. Originally known as *Nurr1*, *Nr4a2* has been implicated in the regulation of *Pomc* expression in the pituitary corticotropic cell line AtT20(Kovalovsky et al., 2002). Furthermore, *Nr4a2* is a pioneer transcriptional regulator important for the differentiation and maintenance of meso-diencephalic dopaminergic (mdDA) neurons during development(Perlmann & Wallen-Mackenzie, 2004). It is crucial for expression of a set of genes including *Slc6a3* (plasma membrane dopamine transporter, DAT), *Slc18a2* (synaptic vesicular monoamine transporter, VMAT), *Th* (Tyrosine hydroxylase), *Foxa1* and *Foxa2* (forkhead family TFs) and *Drd2* (dopamine receptor 2) that together define the dopaminergic neuronal phenotype(Hong et al., 2014; Jankovic, Chen, & Le, 2005; Saucedo-Cardenas et al., 1998; Smits, Ponnio, Conneely, Burbach, & Smidt, 2003). An additional key factor involved in the generation and maintenance of mdDA neurons, *Lmx1a* (Lim- and homeodomain factor 1a) (Andersson et al., 2006; Hong et al., 2014), was also a key feature gene of clusters 3 and 8 (Supplemental Figure 4A and 4D) suggesting that POMC and DA neurons share at least part of their genetic differentiation programs.

Similar to cluster 2, reclustering of the differentially expressed genes from cluster 3 revealed two subclusters (Supplemental Figure 4C). Subcluster 3-1 constituted 72% of the main cluster 3 and was characterized by *Pomc^low^* neurons with strong *Nr4a2* expression at all developmental ages. Feature genes for subcluster 3-1 also included *Lmx1a, Slc6a3, Slc18a2*, *Foxa2* and *Th,* all feature genes for developmental subclusters E13.5-10 and E15.5-11 derived from cluster 3 (Supplemental Figure 7) indicating that a fraction of cluster 3 neurons were dopaminergic and existed transiently during a three-day developmental window (Supplemental Tables 2, 5 and 9). In contrast, subcluster 3-2 neurons were *POMC^high^* together with characteristic expression of the known *Pomc* transcriptional activator genes and were present only at ages P5 and P12. The dissociation of high *Pomc* expression with low *Nr4a2* expression in subcluster 3-2 suggests that the latter is not necessary for hypothalamic *Pomc* expression, unlike its role in pituitary corticotrophs(Murphy & Conneely, 1997).

#### Cluster 8

*Pomc^low^/Pitx2* is more closely related to clusters 2. *Pomc^med^/Ebf1* and 3. *Pomc^med^/Nr4a2* than to any of the others (Supplemental Figure 4D). In addition to *Pitx2,* major feature genes were *Barhl1, Foxa1, Foxp2, Lmx1a/b,* and *Tac1* (Supplemental Table 2). Intriguingly, virtually all of the top cluster 8 defining genes (Supplemental Table 2) are identical to those that characterize both the mesencephalic dopamine neurons and glutamatergic, non-dopaminergic, subthalamic nucleus neurons(Kee et al., 2017). Dual RNAscope in situ hybridization for *Pitx2* and *Pomc* showed limited overlap at E12.5 (Supplemental Figure 4E). Cluster 8 is the only cluster with the neuropeptide *Cck* as a feature gene at developmental ages P5 and P12 (Supplemental Figure S9).

#### Cluster 4

*Pomc^low^/Otp* cells uniquely expressed *Otp* at all developmental ages (Figure 3A). *Pomc* expression was moderate at E11.5 and then dropped to consistently lower levels through age P12 despite continued expression of *Isl1, Nkx2-1* and *Tbx3.* Interestingly, expression of the other transcriptional modulator important for *Pomc* expression, *Prdm12*, was detected only at E11.5. Other characteristic features of cluster 4 were increasing gradients of *Agrp, Npy* and *Calcr* expression over the course of development (Figure 3A). Minimal co-expression of *Pomc* and *Npy* was confirmed by FISH at age E15.5 (Figure 3C). *Sst* was highly expressed in this cluster at ages E11.5, E13.5, P5 and P12 with a temporary drop off at E15.5 and E17.5. Altogether, these patterns of gene expression indicate that a subpopulation of *Pomc* neuronal progenitors differentiates into the canonical AGRP/NPY neurons located in the adult Arc as suggested previously by studies based on transgenic lineage tracing(Padilla et al., 2010). Most of the top enriched genes of this cluster are expressed in the middle and end of pseudotime prediction emphasizing the functions of regulation of inhibitory synapses, GABA-ergic synapses, ribosomal biogenesis, synaptic vesicle cytoskeletal transport and others (Figure 3B). A map of gene expression in individual cells within the cluster 4 UMAP plot demonstrated that *Npy* was expressed in virtually all *Agrp* neurons, but few *Sst* neurons. *Sst* and *Agrp* neurons were essentially separate cell populations (Figure 3D). The GABAergic marker *Slc32a1* encoding the vesicular inhibitory amino acid transporter was expressed at the second highest level in cluster 4 relative to the other clusters (Supplemental Table 1), consistent with the GABAergic phenotype of mature AGRP/NPY neurons (Supplementary Figure 6).

**Figure 3.**
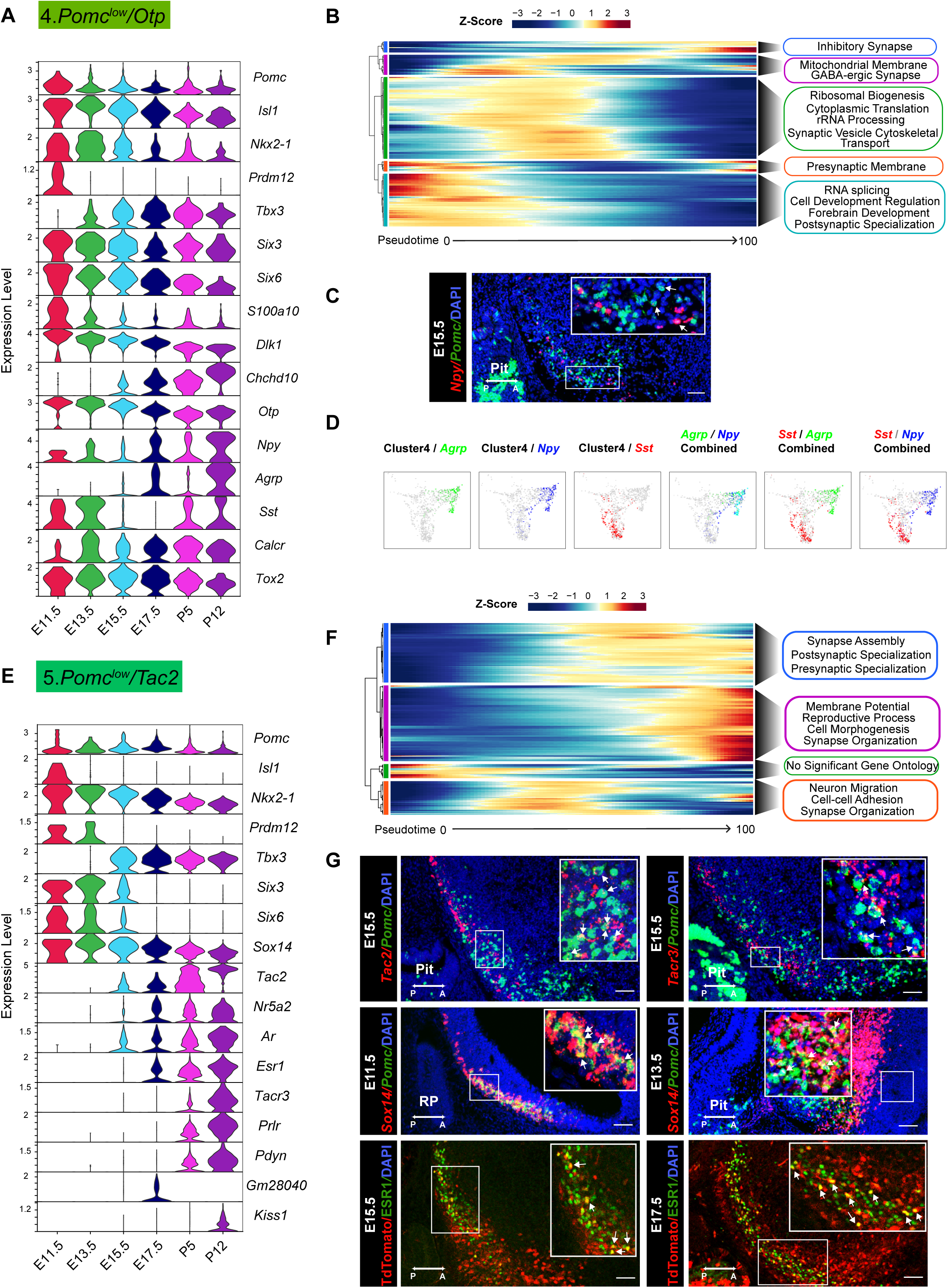
Temporal gene expression patterns of the *Pomc^low^/Otp* cluster and *Pomc^low^/Tac2* cluster. (A) and (E) Violin plots showing the expression of signature genes at each developmental stage in *Pomc^low^/Otp* cluster and *Pomc^low^/Tac2* cluster respectively. (B) and (F) Heatmaps showing the gene ontology analysis based on the top marker genes in the order of pseudotime in *Pomc^low^/Otp* cluster and *Pomc^low^/Tac2* cluster respectively. (C) Fluorescence *in situ* hybridization showing the co-localization of *Pomc* (green) and *Npy* (red) at E15.5. (D) UMAP plots showing cells in the *Pomc^low^/Otp* cluster expressing *Sst*, *Agrp*, *Npy*, *Sst*/*Agrp*, *Sst*/ *Npy* or *Agrp/Npy.* Cells expressing *Sst* and cells expressing both *Agrp*/*Npy* are from two distinct cell populations. (G) Fluorescence *in situ* hybridization showing the co-localization of *Pomc* (green) and *Tac2* (red), the co-localization of *Pomc* (green) and *Tacr3* (red), and the co-localization of *Pomc* (green) and *Sox14* (red) at the indicated developmental stages. Immunofluorescence showing the co-localization of TDTOMATO (red) and ESR1 (green) at the indicated developmental stages. Insets are magnified views of the indicated boxes. Arrows indicate co-expressing neurons in the merged panels. Scale bar: 50 µm. Image orientation: left, posterior; right, anterior; Pit: pituitary gland, RP: Rathke’s pouch.

#### Cluster 5

*Pomc^low^/Tac2* represents a distinct neuronal population from cluster 1 in its patterns of TFs associated with the development of mature POMC neurons (Figure 3E). In particular, *Isl1* was highly expressed only at E11.5 and *Prdm12* at E11.5 and E13.5 in cluster 5. Starting from E15.5 through P12 there was a gradual onset of expression for a set of genes including *Tac2, Ar, Esr1, Tacr3, Prlr, Pdyn,* and *Kiss1/Gm28040* isoforms (Figure 3E). Most of the top enriched genes for this cluster are inferred to be expressed at the middle to end along the pseudotime prediction, and their functions are related to reproductive process, synapse assembly/organization, pre/post synaptic specialization, and neuron migration (Figure 3F). Confirmations of co-expression for *Pomc* with either *Tac2, Tacr3* and *Sox14* or TDTOMATO with ESR1 at several embryonic ages are shown in Figure 3G. Interestingly, *Sox14* has been reported to be necessary for expression of *Kiss1*(Huisman et al., 2019). Taken together, this ensemble of genes is characteristic of adult Arc KNDY neurons that play an essential role in the control of reproductive physiology. A previous publication based on transgene lineage tracing also concluded that a significant subpopulation of KNDY neurons is derived from POMC progenitor cells(Sanz et al., 2015).

A prominent and unique characteristic of **cluster 6.** *Pomc^low/Dlx5^* neurons was the expression of members of the *Dlx* family of homeobox TFs (Supplementary Figure 5A and 5B). The six family members are usually expressed in 3 linked pairs of isoforms: 1 and 2, 3 and 4 or 5 and 6. Cluster 6 was enriched in *Dlx1/2,* and *Dlx5/6* together with *Dlxos1*. The *Dlx1/2* pair is known to activate expression of *Ghrh* in the mouse Arc(Lee et al., 2018). However, there were no transcripts for *Ghrh* expressed in cluster 6 or any of the other *Pomc* neuronal clusters suggesting that *Dlx1/2* are not sufficient to induce *Ghrh* expression. Moreover, it has been recently shown that *Dlx1/2* binding to *Otp* regulatory elements inhibits OTP production, subsequently reducing downstream *Agrp* expression(Lee et al., 2018). Cluster 6 had the highest expression of the opioid prepronociceptin (*Pnoc*) compared to the other integrated clusters, but low levels of preprodynorphin (*Pdyn*), similar to cluster 5 KNDY neurons.

**Cluster 6** had the highest *Gad1*, *Gad2* and *Slc32a1* (*Vgat,* vesicular inhibitory amino acid transporter) expression density of all clusters with virtually no *Slc17a6* (*Vglut2,* vesicular glutamate transporter 2) expression, suggesting that these differentiating neurons were exclusively GABAergic. A comparison of GABAergic and glutamatergic markers among the eight neuronal clusters is shown in Supplementary Figure 6. Cluster 4 neurons were also uniformly GABAergic. Cluster 1 neurons contained a mixture of glutamatergic and GABAergic markers with the exception of undetectable *Slc32a1.* However, they did express low levels of *Slc18a2* (*Vmat2*, vesicular monoamine transporter 2), which possibly functions as an alternative vesicular GABA transporter in some dopamine neurons(German, Baladi, McFadden, Hanson, & Fleckenstein, 2015). This combination of mixed glutamatergic and GABAergic features in cluster 1 is characteristic of postnatal POMC neurons(Jones et al., 2019; Wittmann, Hrabovszky, & Lechan, 2013) and was also present to a limited extent in cluster 2. The remaining neuronal clusters 3, 5, 7 and 8 were all uniformly glutamatergic.

#### Cluster 7

*Pomc^low^/Prdm13* contained the smallest number of cells among the eight neuronal clusters, and the cells were primarily limited to developmental ages E13.5 to E17.5 (Figure 1B). Feature genes included *Prdm13, Nr5a1, Adcypap1, Cnr1, Fam19a1, Rbp1* and *Pdyn* (Supplementary Figure 5C). *Prdm13* was reported recently(X. Chen et al., 2020) to also be highly expressed in E15.5 POMC cells.

We further analyzed all of the neurons and non-neuronal cells by reclustering them based on their individual developmental ages (Supplemental Figure 7) rather than for their gene expression profiles integrated across all ages. Average gene expression levels for each of these developmental subclusters are listed in Supplemental Table 8 and the feature genes defining the developmental subclusters are listed in Supplemental Table 9. Based on these data, it was then possible to trace the temporal continuity between developmental subclusters from age E11.5 to P12 relative to the original integrated cell cluster identities (Supplemental Figure 8). These data are important because they connected the dynamic nature of gene expression profiles at each embryonic stage leading to the eventual transcriptomes present in the postnatal time points.

### Comparison of transcriptional profiles for neuropeptides, neuropeptide processing enzymes, neuroendocrine secretory proteins and GPCRs in the eight neuronal POMC clusters at each developmental age

We identified 19 unique neuropeptide prohormone genes that were clearly differentially expressed features in at least one of the neuronal clusters at one or more developmental ages (Supplemental Figure 9, Supplemental Table 5). In addition to the neuropeptide genes already mentioned as characteristic features of certain clusters, additional neuropeptides were featured in other clusters primarily at postnatal ages P5 and P12. Notable examples are *Gal, Tac1, Adcyap1, Cartpt* and *Pthlh.* Unlike the mosaic of neuropeptide gene expression profiles across clusters and developmental ages, the neuroendocrine secretory proteins, characteristic of dense core granules,were expressed ubiquitously in all eight neuronal clusters with similar gradients across all developmental ages (Supplemental Figure S9). These included members of the chromogranin (*Chga* and *Chgb*) and secretogranin (*Scg2, Scg3* and *Scg5*) families. Among the prohormone processing enzymes, *Pcsk1* was prominently expressed only at P5 and P12 in most neuronal clusters while *Pcsk2* was expressed at both embryonic and postnatal ages in the same clusters as *Pcsk1*. *Pam* was expressed in the same cluster/developmental age pattern as *Pcsk1*. Only *Cpe* was expressed in virtually all neurons at all developmental ages.

Split plots of GPCR gene expression revealed a wide range of cluster- and age-specific patterns (Supplemental Figures 10A and 10B). *Cnr1* (Cannabinoid receptor 1) was an important feature primarily of clusters 7 and the two interrelated clusters 3 and 8. The two metabotropic GABA receptors *Gabbr1* and *Gabbr2* were strongly expressed in all eight neuronal clusters but differed in their developmental gradients. *Gabbr1* was expressed at all developmental ages while *Gabbr2* transcripts were largely present only in more mature neurons at postnatal ages P5 and P12. The opioid receptor *Oprl1* that is selectively activated by nociception/orphanin FQ(Toll, Bruchas, Calo, Cox, & Zaveri, 2016) stood out for its consistently strong expression in all eight neuronal clusters, particularly at postnatal days P5 and P12.

The prokineticin receptor 1 (*Prokr1*) was selectively expressed together with the GPCR accessory protein gene *Mrap2* in cluster 1 *Pomc^high^/Prdm12* neurons. Although MRAP2 was initially identified as an activating modulator of melanocortin receptor signaling, consequently reducing food intake, it was subsequently shown to be promiscuous in its interaction with additional GPCRs(Srisai et al., 2017). Unlike its action on MC4R signaling, MRAP2 inhibits PROKR1 signaling to promote food intake in mice independently of its interaction with MC4R(Chaly, Srisai, Gardner, & Sebag, 2016). Therefore, the co-expression of *Prokr1* and *Mrap2* in anorexigenic POMC neurons suggests an additional mechanism for PROKR1’s regulation of energy homeostasis by modulating the release of melanocortins from POMC neurons. Finally, the high expression of *Npy2r* (neuropeptide Y receptor 2) in both clusters 1 and 4 matches well with its expression in adult anorexigenic POMC neurons and orexigenic AGRP/NPY neurons, respectively.

### TRAP-seq analysis of Pomc-expressing cells at P12 and P60

To define the important genetic programs that direct the transition from early postnatal to adulthood and to compare the differences of the transcriptional programs that guide embryonic vs. postnatal / adult development, we performed Translating ribosome affinity purification TRAP-seq using compound *Pomc-CreERT2; Rosa26-eGFPL10a* transgenic mice where POMC neurons are labeled with a *eGFPL10a* tag after tamoxifen administration at specific developmental stages. Colocalization of POMC and GFP immunoreactivity confirms the expression of *eGFPL10a* in Arc POMC neurons upon tamoxifen injection (Figure 4A). Anti-GFP-conjugated beads were used to pull-down actively translating RNAs bound to the eGFPL10a ribosomes from hypothalamic extracts. Both the pull-down mRNA samples and the resultant supernatant mRNA samples were subjected to RNA sequencing. Principal component analysis (PCA) shows distinct separation of pull-down and supernatant samples at both ages P12 and P60 from three independent experiments, suggesting good quality of these samples and the intrinsic differences between pull-down and supernatant samples (Figure 4B). Both *Pomc* and *GFP* were highly expressed in TRAP pull-down samples relative to the supernatants (Figure 4C), further validating the specificity of the TRAP-seq method. Gene enrichment analysis identified 1143 and 1047 highly enriched genes (*P*<0.05) from *Pomc*-*eGFPL10a* P12 and P60 TRAP-seq pull-downs, respectively (Figure 4D, Supplemental Tables 10 and 11), from which 653 genes expressed in both datasets (Figure 4E). We hypothesize that the commonly expressed genes may be associated with the maintenance of POMC functional identity, whereas the uniquely expressed genes at P12 or P60 may be related to distinct age-dependent biological phenomena. To test this hypothesis, we performed gene ontology analysis on the 653 co-expressed genes, 394 P60 specifically expressed genes and 490 P12 specifically expressed genes. The results demonstrate that the commonly expressed genes are associated with the establishment and maintenance of neuron structure and basic neuronal functions. The top 3 functional annotations include the organization of synapses, regulation of neuron differentiation and cytoskeleton-dependent intracellular transport. The ontology annotations on the uniquely expressed genes from P12 and P60 are dramatically different from each other. The functions of P12 uniquely enriched genes emphasize basic cellular metabolism, cellular modeling and biochemical changes, whereas, the P60 enriched genes are related to the maturation of POMC neurons such as neurotransmitter development and POMC specific function establishment including the responses to nutrient, and the construction of feeding behavioral circuits (Figure 4F). We next compared the TRAP-seq commonly expressed genes to our developmental single cell RNA-seq data to examine their expression levels in each cluster and each age. Approximately one third (241) of the genes shown in the heatmaps on the left are abundantly expressing in the single cell RNA-seq dataset across all the clusters. Notably, compared to the four embryonic stages, a constantly higher transcript abundance was observed at both P5 and P12 ages (grey outlined boxes in Figure 4G), indicating great genetic similarities between P5 and P12. Intriguingly, the patterns of expression levels are very similar in both the P12 and P60 TRAP-seq datasets (Figure 4G, right heatmaps). To gain a better visualization of the distribution of TRAP-seq common genes expression at a single cell level, we grouped the 241 genes as one module, calculated module scores based on the normalized counts and projected the scores to a UMAP plot of the eight neuronal clusters (Figure 4H). Most cells from cluster 1 (*Pomc^high^/Prdm12*), cluster 5 (*Pomc^low/^Tac2*), cluster 3 (*Pomc^med^/Nr4a2*) and a subcluster of cluster 2 (*Pomc^med^/Ebf1*) present a higher module score (high gene expression levels). Assessment together with Figure 5B confirms that most of the high scoring cells are derived from the two postnatal stages (P5 and P12).

**Figure 4.**
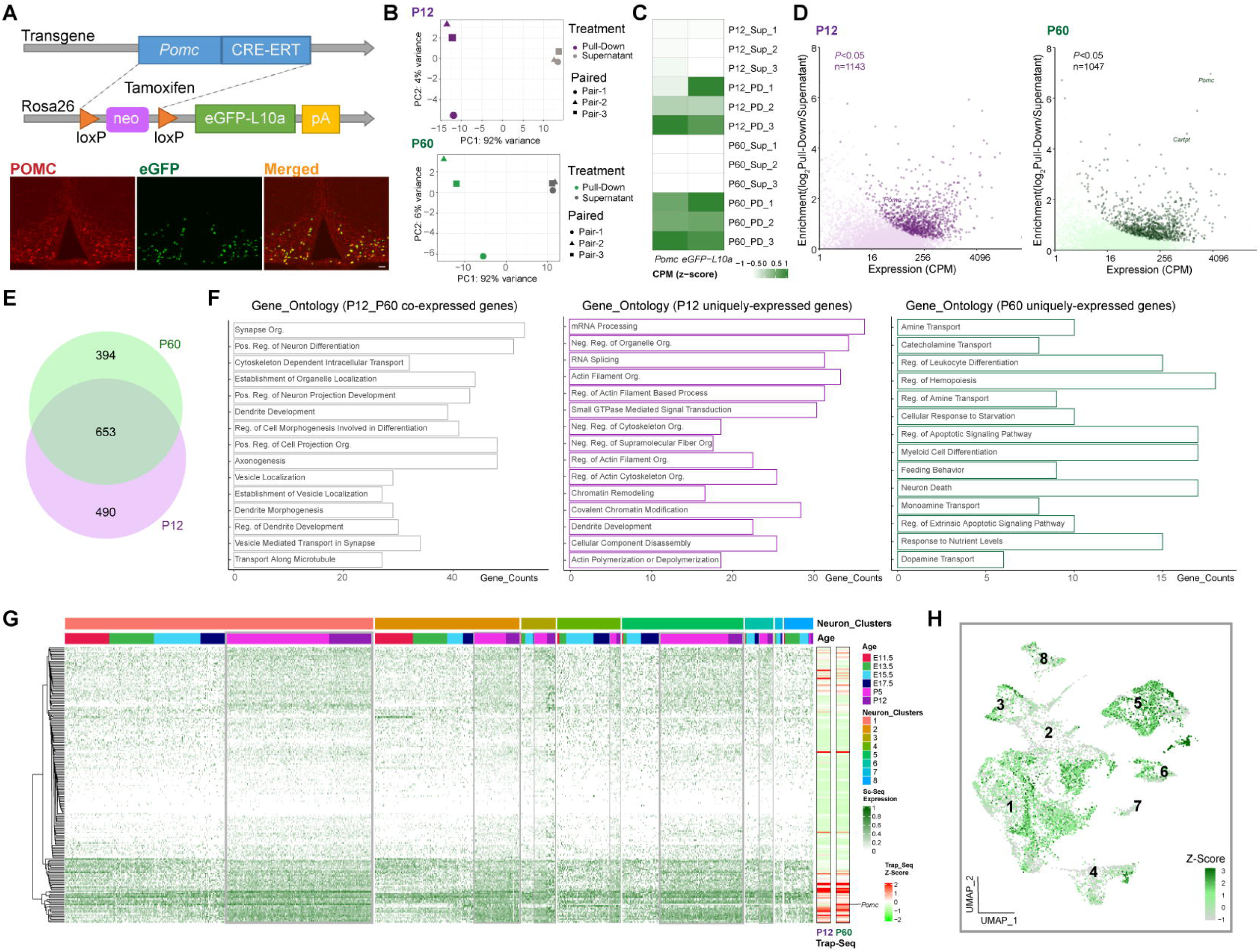
TRAP-seq showing gene enrichment and gene expression profiles at P12 and P60. (A) Schematic diagram showing the generation of *Pomc-CreERT2; ROSA26^eGFP-L10a^* mice for TRAP-seq experiment. Immunofluorescence validated the co-localization of POMC and eGFP. (B) Principal Component Analysis showing the separation of the RNA sequencing data from beads pull-down vs. supernatant at P12 (Purple) and P60 (Green) respectively. (C) Heatmap showing both *Pomc* and eGFP were highly expressed in beads pull-down samples. Rows are the biological replicates of each sample. Sup: Supernatant samples; PD: Pull-Down samples. Data were presented as scaled counts per million (CPM). (D) Gene enrichment plots showing 1143 genes and 1047 genes were significantly expressed in beads pull-down samples at P12 and P60 respectively (*P*<0.05). (E) Venn diagram showing the number of genes highly enriched in both P12 (Purple) and P60 (Green) beads pull-down samples. (F) Gene ontology analysis showing the top 15 biological processes that were represented in P12 and P60 co-expressed genes, P12 uniquely expressed genes and P60 uniquely expressed genes. (G) Expression profile of the top enriched genes from both P12 and P60 TRAP-seq datasets across 8 neuronal clusters; grey boxes indicate the higher expression of these genes in the two postnatal stages. The two heatmaps (right) indicating the top genes expression in P12 and P60 TRAP-seq datasets. (H) UMAP plot showing the distribution of the top enriched genes from both P12 and P60 Trap-seq datasets in 8 neuronal cell clusters. CPM: counts per million; org.: organization; reg.: regulation; pos.:positive; neg.: negative.

**Figure 5.**
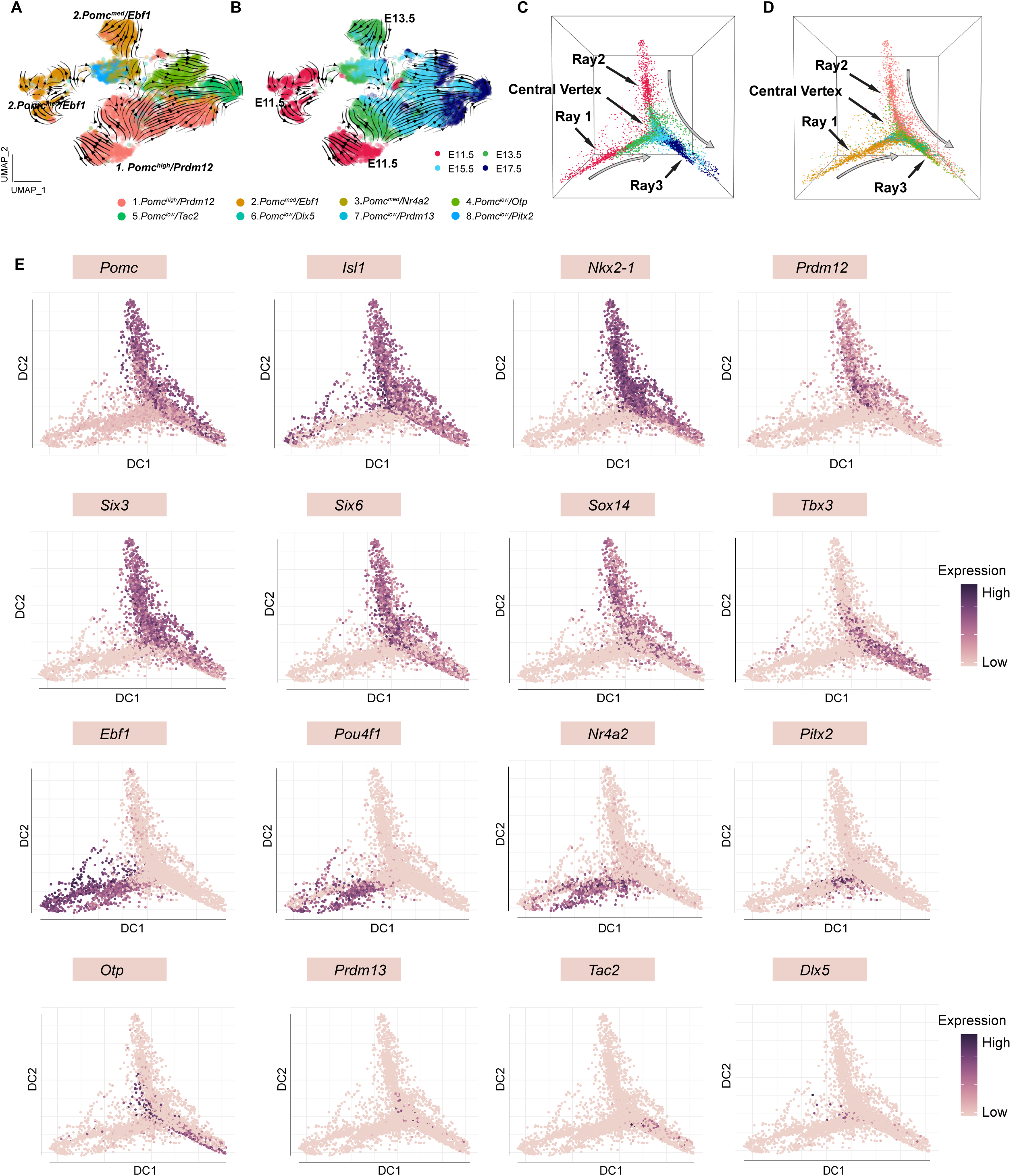
RNA velocity and diffusion maps analyses illustrating the developmental trajectories of hypothalamic neuronal clusters. (A) RNA velocity analysis showing the multiple origins of embryonic POMC cells in 8 neuronal clusters according to Seurat analysis cell embedding. (B) RNA velocity analysis showing the multiple origins of POMC cells correspond to the early embryonic stages. (C-D) Diffusion map showing the cell lineages of POMC neurons during embryogenesis. Cells are colored on the basis of their developmental stage (C) or based on their previously defined clusters (D). (E) Diffusion maps showing the expression of selected genes according to previous characterization of the feature genes representing each cluster. The color intensity corresponds to the normalized gene expression from Seurat analysis.

### Developmental relationships of gene expression patterns in the eight neuronal POMC clusters at four embryonic ages

To investigate the origins and trajectories of POMC progenitors, we initially performed RNA velocity analysis on cells from the 8 neuronal clusters. Since the cells from P5 and P12 are distantly segregated from the four embryonic stages, which subsequently induces noise for cell trajectory analysis, we decide to focus on analyzing cells from the four embryonic stages. Appling cell cluster and age information to RNA velocity analysis reveals multiple progenitor origins including a major origin from the *Pomc^high^/Prdm12* cluster and two minor origins from distinct portions of the *Pomc^med^/Ebf1* cluster (Figure 5A and 5B), which is consistent with the early appearance of these two clusters at E11.5 (Supplemental Figure 11A). Further investigation to cells from E13.5, E15.5 and E17.5 suggest dual origins from *Pomc^high^/Prdm12* or *Pomc^med^/Ebf1* clusters (Supplemental Figure 11B), indicating that a subset of *Pomc*- expressing cells may lose *Pomc* properties during development, which is in agreement with a previous study (Padilla et al., 2010).

To further validate the hypothesis of multiple origins of POMC progenitors and to better visualize cell trajectories in three dimensional Euclidian space, we constructed diffusion maps of all neurons from the four embryonic stages (Figure 5C). The image depicts a triangular beveled star polygon with rays 1 and 2 derived primarily from distinct groups of neurons at ages E11.5 and E13.5, a center vertex containing individual neurons from the intermediate ages E13.5 and E15.5 and a third ray with neurons from ages E15.5 and E17.5. The same data are alternatively color-coded for each neuron’s cluster identity (Figure 5D) showing that ray 1 is primarily derived from cluster 2, ray 2 from cluster 1, the center vertex from a convergence of all clusters 1-8 and ray 3 from clusters 1, 2, 4, 5 and 6. Compared to Ray 2, cells in Ray 1 are not continuously ordered, but instead show an abrupt transition at the junction of cells from E11.5 and E13.5. The diffusion maps analysis appears to confirm the multiple origins of POMC progenitor cells demonstrated by the RNA velocity study.

We then overlaid the geometric location of each neuron with the expression levels of selected feature genes from each of the eight clusters (Figure 5E). Consistent with the diffusion maps analyses, the more granular data for individual genes and neurons illustrates the essentially identical patterns of high *Pomc* expression with the two originally discovered cognate transactivating factors *Isl1* and *Nkx2-1* across embryonic development (Rays 2 and 3). However, *Isl1* was also expressed in numerous *Pomc^low^* neurons at E11.5 and E13.5 (Ray 1). *Prdm12* and *Tbx3* exhibited complementary temporal patterns of high expression at early (Ray 2) versus late (Ray 3) embryonic ages, respectively. The combined developmental and geometric patterns of *Six3, Six6* and *Sox14* expression were all very similar to both *Pomc* and *Nkx2-1*, suggesting that all four of these TF genes might be necessary to define the cell fate of a subpopulation of hypothalamic neuronal progenitors into mature differentiated POMC neurons.

In contrast to those genes with expression patterns that defined the continuum of neuronal identity from Ray 2 to Ray 3, another set of genes including *Ebf1, Pou4f1* and *Nr4a2* was similarly highly expressed only at the two earliest embryonic ages in neurons constituting Ray 1. The expression of *Pitx2,* which characterizes cluster 8 and a portion of cluster 3, was high in Ray 1 as it merges into the center vertex and was mostly derived from age E13.5. Finally, the temporal and geometric patterns of high expression levels for *Otp, Prdm13, Tac2* and *Dlx5* were distinct from each other and the aforementioned genes, consistent with their ontogeny into the distinct differentiated cell fates of cluster 4. AGRP/NPY/GABA, cluster 5. KNDY and cluster 6. GABA/PNOC neurons, respectively.

### Projection of phenotypically distinct cell clusters identified by developmental transcriptomic profiles to neuronal clusters in the adult Arc

We next asked how the transcriptomes of developing POMC neuron progenitors were related to those of neurons in the adult Arc at age P60. The raw single cell sequencing data from Campbell, et al.(Campbell et al., 2017) was transformed, filtered and then combined with our data using the standard data sets integration method from the Seurat package as detailed in the Materials and Methods. Each of the eight *Pomc* neuronal clusters defined from our scRNAseq data and the 34 neuronal clusters from the adult study of Campbell *et al*. (2017)(Campbell et al., 2017) were then projected onto combined UMAP plots to predict the cellular relationships between the two cluster sets (Supplemental Figure 12). A summary of the embryonic to adult cluster projections is shown in Supplemental Figure 13.

66% of cells in the *Pomc^high^/Prdm12* neuronal cluster projected to cluster n14 *Pomc/Ttr* from the adult Arc and another 15% projected to n15 *Pomc/Anxa2* neurons. Although, neither of the adult feature genes *Ttr* or *Anxa2* was expressed significantly in any of the integrated developmental clusters, other key feature genes such as *Pomc, Nkx2-1, Tbx3, Tox2, Six3, Six6, Sox2, and Dlk1* do correlate. All together, these data indicate the continued phenotypic evolution of POMC neurons between ages P12 to P60.

93% of *Pomc^low^/Tac2* cells projected to the adult cluster n20 *Kiss1/Tac2*, providing further evidence that at least a subpopulation of embryonic POMC progenitors further differentiate into mature KNDY neurons. Similarly, there was a 67% correspondence of *Pomc^low^/Otp* neurons with cluster n13 *Agrp/GM8773* neurons, while another 27% of the former projected to n23 *Sst/Unc13c* neurons. These two pairings are consistent with the developmental subclustering of *Pomc^low^/Otp* neurons into those that co-express *Agrp* and *Npy* and those that highly express *Sst* but neither of the other two neuropeptides (Figure 3D). 72% of *Pomc^low^/Dlx5* neurons projected to n22 *Tmem215* neurons. *Tmem215* was identified as a feature gene in combination with *Dlx5* in subcluster P12-12 from our data set (Supplemental Figure 7 and Supplemental Table 9).

78% of cluster 7 *Pomc^low^/Prdm13* neurons projected to the adult neuronal cluster n29 *Nr5a1/Adcyap1*. In the adult hypothalamus, expression of these two genes is characteristic of the ventromedial hypothalamic nucleus instead of the Arc. Expression of the trio of *Prdm13*, *Nr5a1* and *Adcyap1* genes was a significant defining feature unique to cluster 7 *Pomc^low^/Prdm13* neurons from our developmental data set, although *Nr5a1* and *Adcyap1* individually were also feature genes in additional clusters (Supplemental Table 2). Interestingly, the other 11% of *Pomc^low^/Prdm13* neurons projected to cluster n20 *Kiss/Tac2.* Cluster 5 KNDY neurons had significant *Tac1* and *Sox14* expression (Figure 3 and Supplemental Table 2), which may suggest why cluster 7 also projected to the adult n20 *Kiss/Tac2* cluster.

Integrated cluster 2 *Pomc^med^/Ebf1* neurons projected primarily to n34 unassigned2 cells, but 18% projected to n21 *Pomc/Glipr1* neurons and 8% to n14 *Pomc/Ttr* neurons. However, when cluster 2 was reclustered into two distinct components, subcluster 2-1 clearly projected to both clusters n14 *Pomc/Ttr* and n15 *Pomc/Anxa2* neurons, exactly the same as integrated developmental cluster no. 1 *Pomc^high^/Prdm12* canonical POMC neurons. Similarly, integrated cluster 3 *Pomc^med^/Nr4a2* neurons projected mostly to n33 unassigned1 cells but 16% of the neurons projected to n32 *Slc17a6*/*Trhr* neurons. VGLUT2 encoded by *Slc17a6* was most prominently expressed in cluster 3 *Pomc^med^/Nr4a2* neurons compared to all the other integrated clusters.

## Discussion

In this study, we tracked the genetic programs that give rise to *Pomc*-expressing neurons in the developing hypothalamus and followed their progression into terminally differentiated POMC neurons and alternative phenotypic destinies. To this end, we performed a single-cell RNA-seq study specifically of *Pomc*-expressing cells present in developing mouse hypothalami at four embryonic time points (E11.5, E13.5, E15.5, E17.5) and two early postnatal days (P5 and P12). We also performed a complementary TRAP-seq study on affinity purified translating RNAs derived from hypothalamic POMC cells obtained at ages P12 and P60. These developmental stages were selected based on the ontogeny of *Pomc* expression in the presumptive hypothalamus(Japon, Rubinstein, & Low, 1994), peak of neurogenesis of POMC neurons(Nasif et al., 2015; Orquera et al., 2019), subsequent developmental maturation(Padilla et al., 2010; Sanz et al., 2015) and terminal differentiation(Quarta et al., 2019).

Our study revealed the existence of a major group of POMC neurons (Cluster 1. *Pomc^high/^Prdm12*) that showed by far the highest levels of *Pomc* UMIs relative to all other neuronal clusters at all ages. We believe these developing cells constitute the direct precursors of canonical POMC neurons in the adult hypothalamus. Cluster 1 was characterized by a unique combinatorial set of feature TFs that includes *Isl1*, *Nkx2-1* and *Prdm12*. In the last few years, our laboratories demonstrated that these three TFs are present in neurons that start expressing *Pomc* in the developing arcuate nucleus at E10.5, and their expression continues at later developmental ages and adulthood in hypothalamic POMC neurons. Indeed, we showed that ISL1, NKX2-1 and PRDM12 are absolutely necessary to specify the identity of POMC neurons(Hael et al., 2020; Nasif et al., 2015; Orquera et al., 2019). Moreover, we also demonstrated previously that adult mice lacking any of these three TFs exclusively from POMC cells display greatly reduced levels of hypothalamic *Pomc* mRNA together with increased adiposity and body weight. However, our results also indicated that the sole presence of ISL1, NKX2-1 and PRDM12 at E10.5-E11.5 is insufficient to determine the early identity of hypothalamic POMC neurons, suggesting that the combinatorial set of TFs necessary for neuronal-specific *Pomc* expression has additional early components, yet to be discovered. In this regard, our scRNA-seq study revealed several novel candidates in cluster 1 cells including *Six3*, *Six6, Sox1/2/3, Sox14* and other TFs (Figure 2). Future molecular and functional genetic studies will be needed to assess their potential contribution to Arc *Pomc* expression and the functioning of POMC neurons.

*Six3* and *Six6* are two evolutionarily related TF genes known to play critical roles in the development of the eyes, forebrain and pituitary(W. Liu & Cvekl, 2017; W. Liu, Lagutin, Swindell, Jamrich, & Oliver, 2010; Suh et al., 2010). The expression patterns of *Six3* and *Six6* highly overlap during early embryogenesis but segregate into different territories during late embryonic development, indicating possible distinct postnatal functions(Geng et al., 2008). Indeed, *Six3* knockout mice die at birth whereas mice lacking *Six6* exhibit decreased numbers of GNRH neurons and impaired fertility(Larder, Clark, Miller, & Mellon, 2011). Although *Six3* and *Six6* are abundantly expressed in Arc POMC neurons across all examined developmental stages (Figure. 2), their participation in the control of *Pomc* expression and the patterning and function of POMC neurons requires further investigation. Regarding Sox genes, a family of TFs known to regulate cell fate by transactivating cell-specific genes, *in silico* analysis of conserved SOX protein binding motifs suggests the potential binding of SOX proteins including SOX1, SOX2, SOX3, and SOX14 to *<u>Pomc</u>* neuronal enhancers nPE1 and nPE2(X. Chen et al., 2020). All four of these *Sox* genes were highly enriched in *Pomc^high^/Prdm12* cluster 1 (Figure 2 and Supplementary Table 2) as well as in *Pomc^low^/Tac2* cluster 5 and *Pomc^low^/Prdm13* cluster 7. Furthermore, mice lacking *Sox14* exhibit a great reduction in KNDY neurons(Huisman et al., 2019).

A particular feature gene present in cluster 1 was *Tbx3*, a T-box TF that has been recently reported to play a key role in Arc *Pomc* expression(Quarta et al., 2019). Unlike *Isl1*, *Nkx2.1* and *Prdm12*, which are all strictly required at the onset of *Pomc* expression (E10.5), *Tbx3* expression commences several days later (E15.5) in the Arc and, therefore, apparently does not participate in the early specification of POMC neurons. However, *Tbx3* plays an important role in the functional maturation of these neurons as a transcriptional booster of arcuate *Pomc* expression. It was recently demonstrated in mice specifically lacking *Tbx3* from POMC neurons that *Pomc* expression was diminished leading to hyperphagia and obesity(Quarta et al., 2019).

The TRAP-seq results add another layer to our understanding of the genetic machinery that leads to mature functional POMC neurons. Genes that are uniquely enriched at P12 are associated with the basic cellular biology process and cell metabolism whereas genes specifically expressed at P60 are involved in the establishment of POMC neuronal functions. The co-expressed genes from TRAP-seq are highly expressed at P5 and P12 in the single cell sequencing datasets, suggesting that the genetic programs that dominate embryonic POMC neuronal development are substantially different from the programs that guide postnatal POMC neuronal differentiation. Although only one week apart, the cells at P5 share more genetic similarities to the cells at P12 rather than the cells at E17.5.

An unanticipated finding of our study is that Arc *Pomc*-expressing neurons originate from multiple distinct genetic pathways. Neuronal precursors in cluster 1 characterized by the coexpression of feature genes *Isl1*, *Nkx2-1*, *Prdm12*, *Six3*, *Six6*, and the *Sox* genes 1, 2, 3 and 14 (Figure 2) express high *Pomc* levels and give rise to the major group of canonical POMC neurons. In contrast, cluster 2 characterized by expression of the feature genes *Ebf1* and *Pou4f1*, the absence of *Prdm12* and low levels of *Pomc,* differ essentially from cluster 1 POMC neurons at developmental ages 11.5 through 15.5, which give rise to the alternative POMC neurons. Further studies are needed to understand the functional significance of this non-canonical population of *Pomc*-expressing neurons that originate from an alternative route.

The identification of *Pomc^low^/Tac2* and *Pomc^low^/Otp* clusters confirmed that some fertility-regulating KNDY neurons(Sanz et al., 2015) and orexigenic AGRP/NPY neurons(Padilla et al., 2010) arise from *Pomc* progenitors. Moreover, our data indicate that *Pomc* precursors may also give rise to four other less characterized neuronal populations noted as *Pomc^med^/Nr4a2*, *Pomc^low^/Pitx2*, *Pomc^low^/Dlx5* and *Pomc^low^/Prdm13*. The *Pomc^med^/Nr4a2* and *Pomc^low^/Pitx2* are related clusters sharing 20% of their top 150 feature genes and both of them projected to the same two heterogeneous clusters n33 and n34 in adult Arc neurons (Supplemental Figure S13). Interestingly, many of the top feature genes in cluster 8 *Pomc^low^/Pitx2*, including *Pitx2, Lmx1a, Nr4a2, Foxa1, Foxp2*, and *Barhl2* were reported previously as marker genes for the supramammillary nucleus (SMN) (Kim et al., 2020). This cluster also had the most abundant *Cck* transcripts, further indicating that it may represent a group of cells from the SMN considering the distribution of *Cck*-expressing neurons in the brain(Burgunder & Young, 1989). Notably, POMC cells are undetectable in the SMN in adult mice, suggesting that early *Pomc* expression in these cells may disappear during postnatal growth. This holds true when we compared our datasets to Lam *et. al’s* study profiling FACS sorted individual POMC-EGFP cells from adult hypothalamus(Lam et al., 2017). None of the clusters from their study were characterized by *Pitx2*, *Lmx1a* or *Foxa1* expression. Similarly, the *Pomc^low^/Prdm13* cluster failed to match any clusters from Lam *et. al’s* study(Lam et al., 2017), which is not unexpected since this cluster had the most abundant *Nr5a1* transcripts and corresponded to the n29 *Nr5a1/Adcyap1* (Supplemental Figure S13) cluster(Campbell et al., 2017), representing a cell population in the ventromedial hypothalamus featured by *Nr5a1* (*Sf1*) expression. Immunohistochemistry showed that NR5A1 is abundantly co-localized in *Pomc*-expressing cells at early embryonic days (E11.5-15.5) but not later perinatal days (E17.5) (Figure 2C). Conversely, cluster 2 adult POMC cells (featured with *Fzd5*, *Setd4* and *4732456N10Rik*) from Lam *et.al’s.* study failed to match any of the clusters in our study, indicating that this cluster may contain cells generated postnatally. The signature genes for our *Pomc^low^/Dlx5* cluster including *Dlx1, Dlx2, Dlx5*, and *Dlx6* were highly enriched and specific for cluster 1 in Lam *et. al’s* study, indicating that the *Pomc^low^/Dlx5* cluster may become a subset of adult POMC neurons. Other than the above four less defined clusters, the *Pomc^high^/Prdm12* cluster is highly similar to cluster 4 in Lam *et. al’s* study characterized by *Prdm12, S100a10, Cpne4* and *Calb1* expression, representing the classic POMC neurons in adult mice.

The *Pomc^med^/Nr4a2* cluster, particularly subcluster 3-1, is the second cluster enriched with *Pitx2, Lmx1a,* and *Foxa1* expression and also has the most abundant *Nr4a2*, *Pitx3*, and *Th* transcripts (Supplemental Table1). The early B cell factors (*Ebf1-3*) were also present in this subgroup of cells in a temporally restricted expression pattern (Figure 2D). Since the *Ebfs* control differentiation or migration of MbDA subpopulation in a temporally-dependent manner(Yin et al., 2009), it is reasonable to infer that *Pomc^med^/Nr4a2* subcluster 3-1 (Supplemental Figure S4C) may define a subpopulation of immature MbDA neurons. Indeed, *Nr4a2* is expressed in 18.5% of cells in cluster 4 of adult POMC cells from the Lam *et.al.* study(Lam et al., 2017).

It is worth mentioning that despite some extent of similarities between embryonic and adult POMC cells, the majority of cells profiled in our study are distantly related to mature adult POMC neurons. For example, most adult POMC cells express proprotein convertase PC1/3 (*Pcsk1*) and leptin receptor (*Lepr*), whereas only 22% and 2.7% of cells from our datasets express the two genes, respectively, indicating dramatic transcriptional signaling changes in POMC neurons during postnatal maturation after age P12.

## Concluding remarks

In summary, this study is the first to comprehensively characterize transcriptomes of Arc hypothalamic POMC cells during embryonic and early postnatal development. This dataset extends our understanding of the diversification of early POMC cells and provides a valuable resource for further elucidating the regulatory mechanism for *Pomc* expression and neuronal maturation. The identification of new marker genes in each subpopulation from single cell RNA-seq and uniquely expressed genes at P12 or P60 from the TRAP-seq study will potentially lead to new directions for future functional studies of POMC neurons.

## Acknowledgements

We thank Dr. Graham L. Jones, Dr. Zoe Thompson and Charles Keane for their assistance in animal dissection, tissue sectioning and data analysis, Dr. Joshua Welch and Tongyu Liu for advice on data analysis, the Flow Cytometry Core, the Advanced Genomics Core, the Bioinformatics Core and the Microscopy, Imaging and Cellular Physiology Core (supported by NIH Grant: P30DK020572) and the University of Michigan Center for Gastrointestinal Research (supported by NIH grant: P30DK034933) at University of Michigan for data collection.

## Author Contributions

H.Y., M.R., and M.J.L. jointly conceived the study, designed the experiments, analyzed data and drafted the manuscript. H.Y. conducted the experiments. All of the authors edited and approved the final manuscript.

## Methods

### Mice

All procedures were performed in accordance with the Institutional Animal Care and Use Committee (IACUC) at the University of Michigan and followed the Public Health Service guidelines for the humane care and use of experimental animals. Mice were housed in ventilated cages under controlled temperature and photoperiod (12-h light/12-h dark cycle, lights on from 6:00 AM to 6:00 PM), with free access to tap water and laboratory chow (5L0D, LabDiet). Breeding mice were fed with the breeder chow diet (5008, LabDiet). Transgenic mice expressing the fluorescent protein Tdimer-Discosoma red (TdDsRed) in POMC neurons were generated previously^17^. The vast majority of POMC-TdDsRed cells were validated as authentic POMC neurons based on co-localization of ACTH immunostaining with the POMC-TdDsRed fluorophore (Hentges, Otero-Corchon, Pennock, King, & Low, 2009) *Pomc-CreERT2* and Rosa26^eGFP-L10a^ (Stock no. 024750, The Jackson Laboratory) mice were generated as previously described (J. Liu et al., 2014). To induce POMC neurons specific expression of eGFP-L10a at P12 or P60, tamoxifen (50 mg/kg) was injected intraperitoneally for five constitutive days from P6 to P10 or P50 to P54.

### Generation of single cell (sc) suspension for scRNAseq

Adult male POMC-TdDsRed/+ mice were bred with female POMC-TdDsRed/+ mice and the day of copulation plug detection was counted as embryonic day 0.5 (E0.5). Embryonic hypothalami were dissected at E11.5, E13.5, E15.5, and E17.5. Postnatal hypothalami were isolated from pups at postnatal day 5 (P5) and P12. In each dissection, hypothalami from at least 6 pups were pooled together to acquire enough cells for fluorescence-activated cell sorting (FACS). Pooled tissue samples were digested in papain solution supplemented with DNAse for 20 mins (E11.5 and E13.5), 30 mins (E15.5), 40 mins (E17.5) or 1hr (P5, and P12) with gentle agitation based on a published protocol^14^. The isolated cells were stained with DAPI as an indicator for cell viability and sorted by FACS for the transgenic red fluorophore (Supplemental Figure 14). The collected cells were counted and subjected to scRNA-seq library preparation and sequencing (10x Genomics).

### Single Cell RNA Sequencing Data processing

A total of 37,053 single cells (E11.5: 5,329, E13.5: 7,051, E15.5: 5,148, E17.5: 2,488, P5: 7,551 and P12: 9,486) from six different developmental stages were processed using the 10X Genomics Chromium system. The libraries were sequenced on Illumina HiSeq 4000 and NovaSeq platforms. We obtained a total of over 2.6 billion reads with an average of 70,7896 reads per cell. Over 85% of reads mapped confidently to the mouse genome across all six developmental stages. Raw reads were processed with Cell Ranger (version 2.2 and version 3.0). Seurat package (version 3.1.5)(Butler, Hoffman, Smibert, Papalexi, & Satija, 2018) was used for downstream analysis. Since the number of genes and UMI counts varied across developmental stage we applied different criteria at each stage to filter out possible doublets and low quality cells. Specifically, at E11.5, we removed outlier cells that had UMI counts < 5,000 or >60,000 and gene counts <1,000 or >8,000 (determined by the visualization of UMI counts and gene count distributions). At E13.5, we removed outlier cells that had UMI counts <2,500 or >30,000 and gene counts >1,500. At E15.5, we removed outlier cells that had UMI counts >40,000 and gene counts <1,000 or > 6,000. At E17.5, we removed outlier cells that had UMI counts >25,000 and gene counts <1.000 or > 5,000. At P5, we removed outlier cells that had UMI counts <1,500 or >60,000 and gene counts <1,500 or >8,000. At P12, we removed outlier cells that had UMI counts<2,000 or >60,000and gene counts <1,500 or > 8000. Moreover, cells with high proportions of mitochondrial genes (>10%) or hemoglobin genes (>10%) were filtered out. Finally, we removed all the cells without any *Pomc* transcript UMI counts. A total of 13,953 cells passed the criteria (E11.5: 1,498, E13.5: 1,796, E15.5: 2,078, E17.5: 1,139, P5: 3,909 and P12: 3,533) for downstream analysis.

The gene expression level for each cell was normalized by the total expression, multiplied by a scaling factor of 10,000 followed by log-transformation(Butler et al., 2018). The top 2,000 most variable genes were selected from each age and 60 integration anchors were used to combine all the datasets together. After scaling and centering the integrated dataset, we performed principal components (PC) analysis on the data matrix and the top 50 PCs were selected based on JackStraw and Elbow plots from Seurat Package^19^ for data visualization using Uniform Manifold Approximation and Projection (UMAP) technique. We next constructed a Shared Nearest Neighbor (SNN) graph by setting an expected number of neighbors to 50. In order to cluster the cells, a SNN modularity optimization technique within a function FindClusters was used to group cells together with a resolution parameter 0.1. The identity for each cluster was assigned based on the level of *Pomc* transcripts and prior knowledge of marker genes. The similar analysis was applied to define *TdDsRed* cell clusters (Supplementary Figure 2 and Supplementary Figure 3). In addition to the above criteria applied to remove unwanted cells, we chose cells with at least one *TdDsRed* transcript UMI count and used the top 2,000 highly variable genes for principal components analysis. The top 30 PCs were selected for unsupervised clustering with a resolution of 0.3.

Unsupervised cell clustering from each developmental age (Supplementary Figure 6) was conducting using the following PC numbers (E11.5: 25; E13.5:24, E15.5: 26, E17.5: 19, P5: 40, P12: 34) with their corresponding resolutions (E11.5: 0.2; E13.5: 0.6, E15.5: 0.5, E17.5: 0.5, P5: 0.5, P12: 0.5. The UMAP plots, violin plots, feature plots and dot plots were all generated in the Seurat Package. The average expression levels for each gene within each developmental subcluster (Supplemental Table 8) were obtained from Seurat and calculated by the average of [*exp(logNormalized counts) -1*]. The differentially expressed genes in each identified cluster was identified by the comparison of gene expression levels in a specific cluster to all the other clusters.

### Cluster identity comparisons

We took the following steps to compare cluster identities from developmental POMC cells to 34 adult ARC neuronal clusters(Campbell et al., 2017) (Supplementary Figure 12 and Supplementary Figure 13). 1) We obtained the adult ARC single cell sequencing data from https://hemberg-lab.github.io/scRNA.seq.datasets/mouse/brain(Kiselev, Yiu, & Hemberg, 2018) and transformed the dataset from SingleCellExperiment format to Seurat object format. 2) We normalized adult ARC data and acquired the top 2000 most variable gene for further analysis. 3) The standard datasets integration method from Seurat package was performed with 50 integration anchors and used to combine the two datasets. 4) The integrated data were scaled and the top 50 PCs were selected for unsupervised clustering implemented in Seurat. 5) We chose 50 transfer anchors to predict the developmental POMC cells’ identities based on adult ARC cell clusters. A similar approach was applied to compare cluster identities between POMC cells and DSRED cells (Supplementary Figure 2) with the top 50 anchors for data integration and 50 anchors to transfer cell identities from POMC cells to DSRED cells. We also applied this approach to conduct age-pairwise comparisons (Supplementary Figure 8) with default setting from Seurat V3.1.5 package.

### Cell lineage construction

For RNA velocity analysis, spliced and unspliced matrices of reads were summarized using velocyto (v.0.17.16) with default parameters (La Manno et al., 2018). Low complexity and repeat regions were downloaded from the UCSC browser (rmsk table from mm10). scVelo was performed for RNA velocity analysis (v.0.2.2) (Bergen, Lange, Peidli, Wolf, & Theis, 2020). The cell embedding information was acquired from Seurat analysis. The top 4000 genes, the top 20 principal components and the top 20 neighbors under stochastic mode were used to generate velocity graph.

Cell lineage was constructed using the Destiny (version 2.14.0) package, which implements the formulation of diffusion maps(Angerer et al., 2016). Diffusion maps are a spectral method for non-linear dimension reduction, which is especially suitable for analyzing single-cell gene expression data from different time-courses. We removed VLMC, glial cells and astrocytes (cluster 9, 10 and 11 in Figure1) and kept only neurons at embryonic stages (E11.5, E13.5, E15.5 and E17.5), resulting in a total of 6,005 cells for this analysis. The log-transformed normalized data from Seurat data slot and the corresponding annotation information were imported to construct an expression matrix. The diffusion maps were constructed under the default setting for the Gaussian kernel width sigma (σ) and 300 nearest neighbors. The first two Diffusion Components (DCs) were used to visualize the results. The 3D plots were produced using rgl package (version 0.100.50) with the top three DCs.

To perform pseudotime gene expression analysis, cells from cluster 1, cluster 2, cluster 4 and cluster 5 were extracted from the Seurat object respectively. Raw counts were acquired to construct Monocle (v.2.4.0)(Qiu, Hill, et al., 2017; Qiu, Mao, et al., 2017; Trapnell et al., 2014) datasets. Cells were ordered along the pseudotime by setting E11.5 as the root state. Differentially expressed genes (Log2FoldChange > 0.25 and *P*<0.05) in each cluster from the Seurat analysis were plotted along the pseudotime using “plot_pseudotime_heatmap” function. Gene ontology analysis (Biological Process) were performed using ClusterProfiler (v.3.18.1) (Yu, Wang, Han, & He, 2012).

### TRAP-seq analysis

Mice (*Pomc-CreERT2*; ROSA26^eGFP-L10a^) were euthanized and decapitated. The brain was removed from the skull and the arcuate nucleus was isolated from 2-mm thick coronal slices using a brain matrix. Tissues were then homogenized and the subsequent tissue lysate was subjected to immunopurification steps according to the previous protocol (Heiman, Kulicke, Fenster, Greengard, & Heintz, 2014). The ribo-depletion kit (RiboGone, Takara, CA) and SMARTer Stranded total RNA sample prep kit were used to remove excess ribosomal RNA and synthesize cDNA library. Samples were sequenced on a 50-cycle single end run on a HiSeq 4000 (Illumina) according to manufacturer’s protocols.

Raw sequencing data was processed at the University of Michigan Bioinformatics core. Briefly, the quality of the raw reads data was checked using FastQC (v.0.11.3) and the filtered reads were aligned to reference genome (UCSC mm10) using TopHat (v.2.0.13) and Bowtie2 (v.2.2.1) with default parameters. The HTSeq/DEseq2 method was used for differential expression analysis with paired-samples and treatment (pull-down vs. supernatant) as the main effects.

### Tissue collection and Immunofluorescence staining

Depending on developmental stage, embryos at E10.5 – E17.5 were fixed in 4% paraformaldehyde in phosphate buffer solution (PBS) at 4 °C for a various period of time. Specifically, embryos at E10.5 to E12.5 were fixed for 1h; heads from embryos at E15.5-E16.6 and E17.5 were fixed for 2 h and 4 h respectively. Tissues were stabilized in 10% sucrose/10% gelatin in PBS at 37 °C for 30 min prior to embedding in O.C.T. compound as described previously^9^. Embryonic brains were sectioned sagittally on a cryostat (Leica CM1950) at 20 μ m thickness. For dual immunofluorescence, after washing excess O.C.T. compound with PBS, a heat-induced antigen retrieval process was performed using citrate buffer (10 mM anhydrous citric acid and 0.05% Tween-20, pH 6.0) at 80 °C for 30 min followed by two washes in PBS. Sections were then blocked with 3% normal goat serum and 0.1% Triton X-100 for 1 hour and incubated with primary antibodies at room temperature overnight. After PBS washes, sections were incubated with goat secondary antibodies for 2 hours at room temperature. Nuclei were stained with DAPI (1 mg/l) for 10 min and the slides washed five times with PBS before mounting with ProLong™ Gold Antifade Mountant (ThermoFisher). Sections were imaged on a Nikon 90i fluorescence microscope with NIS-Elements software. Information on the sources and dilution of antibodies are listed in Supplementary Table 12.

### *In situ* hybridization

Fluorescence *in situ* hybridization (FISH) was performed using RNAscope^®^ Multiplex Fluorescent V2 Assay (Advanced Cell Diagnostics) according to the manufacturer’s instructions with slight modifications. Specifically, after washing with PBS, sections were post-fixed with 4% paraformaldehyde for 20 minutes, and dehydrated in a series of ethanol solutions (50%, 70% and 100%). After drying, dehydrated sections were incubated in RNAscope^®^ Hydrogen Peroxide solution for 10 mins followed by the protease treatment using RNAscope^®^ Protease IV for 20 mins. Sections were then hybridized to target RNA probes for 2 hours in the HybEZ^TM^ II hybridization oven. Hybridized probes were amplified using a cascade of signal amplification solutions (AMP 1-3) followed by standard signal developing protocol as described in the manufacturer’s brochure. Detailed information on RNA probes and dilutions are listed in Supplementary Table10. Confocal images were obtained using a Nikon Instruments A1 Confocal Laser Microscope with NIS-Elements software.

## Supplemental Data

**Figure S1**

Heat map showing the expression of the top 30 marker genes defining each cluster from Figure 1. M: microglia; V: vascular and leptomeningeal cells; A: astrocytes.

**Figure S2**

Analysis of POMC cells with *DsRed* transcripts reveals 9 distinct cell clusters. (A) UMAP plot showing the distribution of major cell clusters. (B) Violin plots showing signature gene expression in each cell type. Three feature genes were chosen to represent each cell type. (C) UMAP plots showing the enrichment of feature genes in each cluster. Genes representative of each cluster were colored and highlighted to show the cluster-specific enrichment. The color intensity corresponds to the normalized gene expression from Seurat analysis. (D) Identification of similar cell clusters between POMC cells with *DsRed* transcripts and POMC cell clusters. Red: POMC cell clusters are from Figure 1. Blue: POMC cells with *DsRed* transcripts from Supplemental Figure 2A. (E) Venn diagram showing the number of cells expressing *Pomc* transcripts, *DsRed* transcripts or both *Pomc* and *DsRed* transcripts. Pie chart showing the percentage of POMC cells expressing *DsRed* transcripts at each developmental stage. (F) Vertical bar-chart showing the percentage of cells with *DsRed* transcripts projected to corresponding POMC cell clusters. VLMC, vascular and leptomeningeal cell.

**Figure S3**

Overview of cell clusters from POMC cells with *DsRed* transcripts. (A) UMAP plot showing the distribution of major cell clusters and their corresponding cell% from each developmental stage. (B) UMAP plots showing the number of cells at each developmental stage. (C) UMAP plots showing the splitting of clusters by developmental stages.

**Figure S4**

Temporal gene expression patterns of the *Pomc^med^/Nr4a2* and *Pomc^low^/Pitx2* clusters. (A) Violin plots showing the expression of signature genes in the *Pomc^med^/Nr4a2* cluster at each developmental stage. (B) Fluorescence *in situ* hybridization showing the co-localization of *Pomc* (green) and *Nr4a2* (red). (C) Re-clustering cells from the *Pomc^med^/Nr4a2* cluster reveals two sub-clusters denoted as 3-1 *Pomc^low^* and 3-2 *Pomc^high^* clusters. Violin plot showing the signature gene expression in 3-1 *Pomc^low^* and 3-2 *Pomc^high^* clusters. (D) Violin plots showing the expression of signature genes at each developmental stage in the *Pomc^low^/Pitx2* cluster. (E) Fluorescence *in situ* hybridization showing the co-localization of *Pomc* (green) and *Pitx2* (red). Scale bar: 50 µm. Insets are magnified views of the indicated boxes. Arrows indicate co-expressing neurons in the merged panels. Scale bar: 50 µm. Image orientation: left, posterior (P); right, anterior (A). RP: rathke’s pouch; Pit: pituitary gland.

**Figure S5**

Temporal gene expression patterns of the *Pomc^low^/Dlx5* and *Pomc^low^*/*Prdm13* clusters. (A) Violin plots showing the expression of signature genes at each developmental stage in the *Pomc^low^/Dlx5* cluster. (B) Fluorescence *in situ* hybridization showing the co-localization of *Pomc* (green) and *Dlx5* (red). (C) Violin plots showing the expression of signature genes at each developmental stage in the *Pomc^low^/Prdm13* cluster. (D) Fluorescence *in situ* hybridization showing the co-localization of *Pomc* (green) and *Prdm13* (red). Insets are magnified views of the indicated boxes. Arrows indicate co-expressing neurons in the merged panels. Scale bar: 50 µm. Image orientation: left, posterior; right, anterior; Pit: pituitary gland.

**Figure S6**

Distribution of GABAergic and glutamatergic neurons in POMC^DsRed^ cell clusters (A) UMAP plots showing the distribution of cells with *Gad1* or *Vglut2* gene expression. (B) Violin plots showing expression of the GABAergic or glutamatergic marker genes in each neuronal cluster.

**Figure S7**

UMAP plots showing the distribution of major cell clusters at each developmental stage. (A) E11.5, (B) E13.5, (C) E15.5, (D) E17.5, (E) P5 and (F) P12. The developmental subclusters derived from each primary cluster are colored to corresponding cells in the UMAP plots.

**Figure S8**

POMCDsRed developmental cell subclusters shown in Supplemental Figure 7 are projected to the original integrated cell clusters from Figure 1. The projections were based on the Seurat analysis and manual comparisons of feature genes in each subcluster (Supplemental Table 2 and supplemental Table5). The clusters from individual stages leading to the same final integrated clusters are labeled with the same color. Question marks indicate uncertainties of individual projections.

**Figure S9**

Overview of neuropeptide, secretory granule and processing enzyme gene expression patterns across clusters and developmental ages. The size of dots indicates the percentage of cells expressing the specific transcripts. The color intensity corresponds to the levels of normalized gene expression.

**Figure S10 A and B**

Overview of G-protein coupled receptor (GPCR) gene expression patterns across clusters and developmental ages. The size of dots indicates the percentage of cells expressing the specific transcripts. The color intensity corresponds to the levels of normalized gene expression. The list of genes was acquired from https://www.guidetopharmacology.org/targets.jsp and Gene Ontology (GO:0005576).

**Figure S11**

**(A)** UMAP plots showing the splitting of neuronal clusters by developmental stages. Clusters were dash-dot circled to show the emerging / evolving progress through development. (B) RNA velocity analysis showing the multiple origins of POMC cells are mainly from 1. *Pomc^high^/Prdm12* and 2. *Pomc^med^/Ebf1* clusters and are mainly from E11.5 and E13.5. (C) RNA velocity analysis on neurons from three late embryonic stages (E13.5, E15.5, and E17.5) showing two origins of POMC cells from 1. *Pomc^high^/Prdm12* and 2. *Pomc^med^/Ebf1* clusters at E13.5.

**Figure S12**

Identification of similar clusters between the neuronal POMC^DsRed^ developmental clusters and the neuronal clusters derived from adult arcuate nucleus(Campbell et al., 2017). (A) UMAP plots showing the integration of POMC^DsRed^ clusters and adult arcuate nucleus neuronal clusters. (B) The percentage of POMC^DsRed^ cells projecting to corresponding adult arcuate nucleus neuronal clusters.

**Figure S13**

Projection of embryonic POMC^DsRed^ cell subtypes to adult arcuate nucleus neuronal clusters. Projected cell clusters from the *Campbell et al.* study ^14^ were labeled with the same color as the POMC^DsRed^ cells clusters. The adult ARC neuronal clusters highlighted in bold font indicate a higher percentage of cells projected to this cluster from the corresponding developmental clusters. The detailed cell embedding and percentage of cells projecting to adult Arc neuronal clusters are shown in Supplemental Figure S12.

**Figure S14**

Fluorescence-activated cell sorting (FACS) strategy for collecting the single POMC-TdDsRed cells used for scRNAseq.

**Table S1 (corresponding to Figure 1)**

**Table S2 (corresponding to Figure 1)**

**Table S3 (corresponding to Supplemental Figure 2)**

**Table S4 (corresponding to Supplemental Figure 3)**

**Table S5 (corresponding to Figures 2 and 3 and Supplementary Figures 4 and 5)**

**Table S6 (corresponding to Figure 2G, cluster 2 and Supplemental Figure 4C, cluster 3)**

**Table S7 (corresponding to Figure 2G, cluster 2 and Supplemental Figure 4C, cluster 3)**

**Table S8 (corresponding to Supplemental Figure 7)**

**Table S9 (corresponding to Supplemental Figure 7)**

**Table S10 (corresponding to Figure 4)**

**Table S11 (corresponding to Figure 4)**

**Table S12 Table of key resources.**

