## Supplementary figures and images for "Developmental single-cell transcriptomics of hypothalamic POMC progenitors reveal the genetic trajectories of multiple neuropeptidergic phenotypes"

### Supplementary Figure1

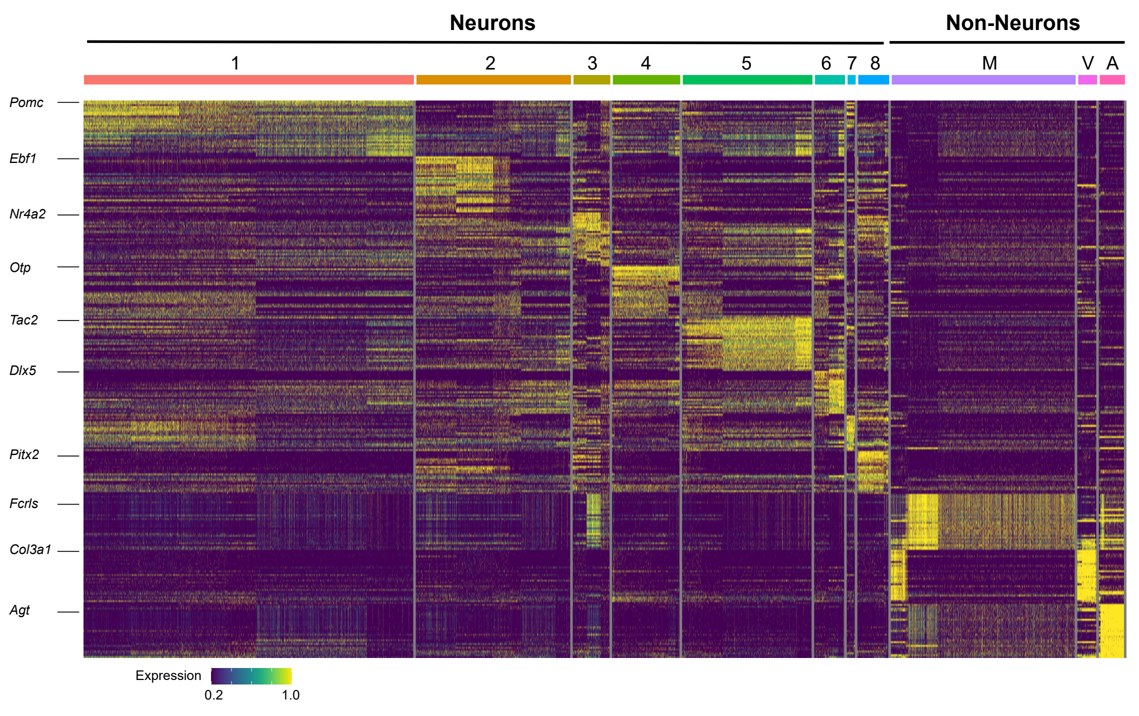

### Supplementary Figure2

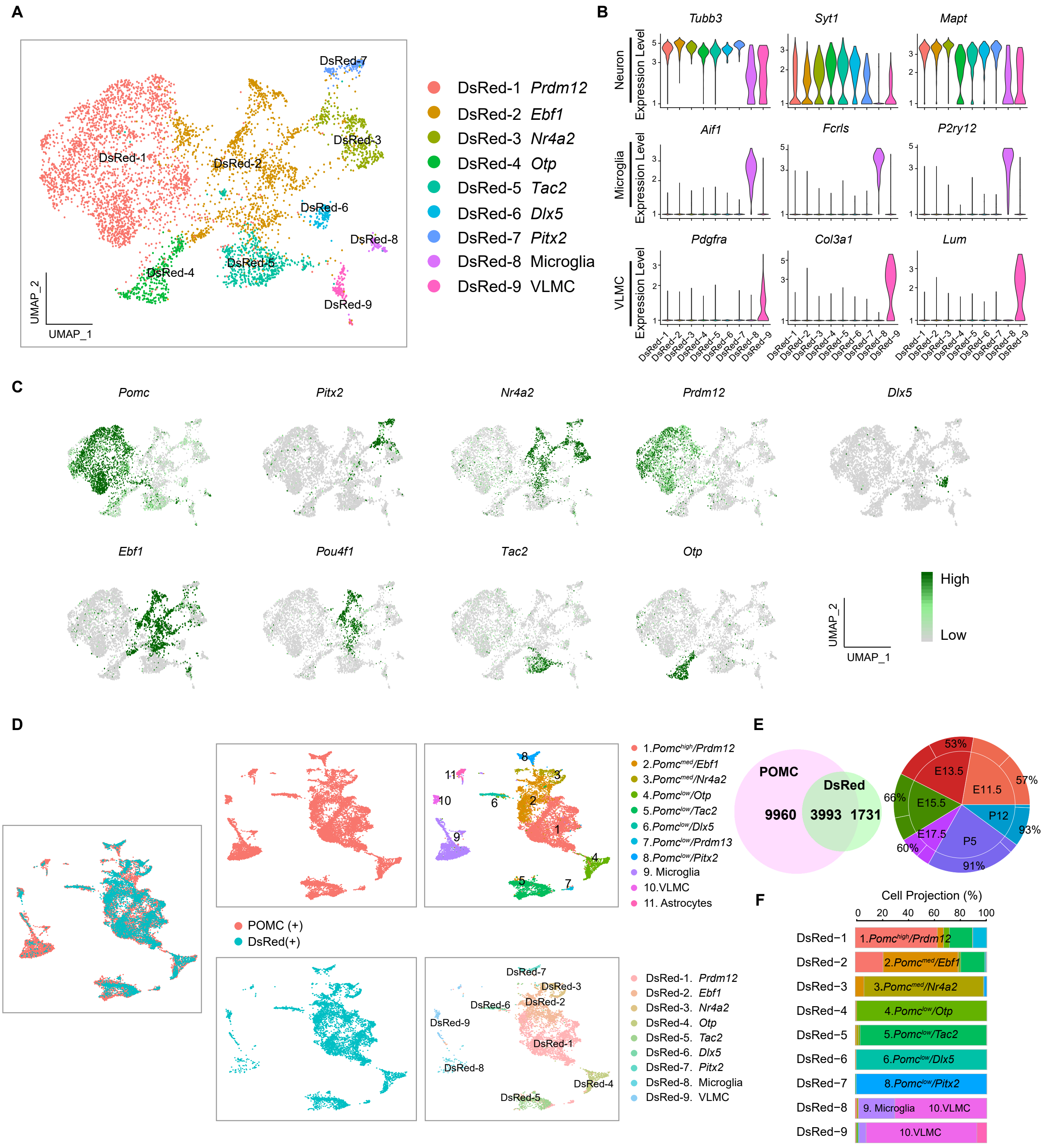

### Supplementary Figure3

**A**

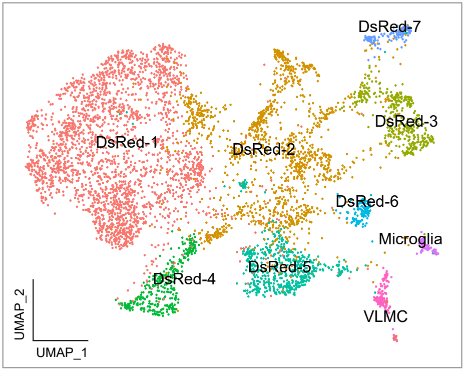


| | | E11.5 | E13.5 | E15.5 | E17.5 | P5 | P12 |
|---|---|---|---|---|---|---|---|
| DsRed-1 | *Prdm12* | 38% | 38% | 40% | 33% | 57% | 69% |
| DsRed-2 | *Ebf1* | 52% | 42% | 7% | 9% | 13% | 15% |
| DsRed-3 | *Nr4a2* | 4% | 5% | 12% | 2% | 12% | 4% |
| DsRed-4 | *Otp* | 2% | 5% | 17% | 14% | 2% | 2% |
| DsRed-5 | *Tac2* | 2% | 3% | 17% | 38% | 11% | 6% |
| DsRed-6 | *Dlx5* | 0% | 1% | 5% | 3% | 4% | 3% |
| DsRed-7 | *Pitx2* | 0% | 6% | 2% | 0% | 1% | 2% |

**B**

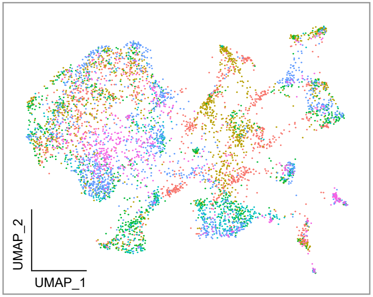


| | Cell # |
|---|---|
| E11.5 | 1273 |
| E13.5 | 1162 |
| E15.5 | 926 |
| E17.5 | 465 |
| P5 | 576 |
| P12 | 1322 |

**C**

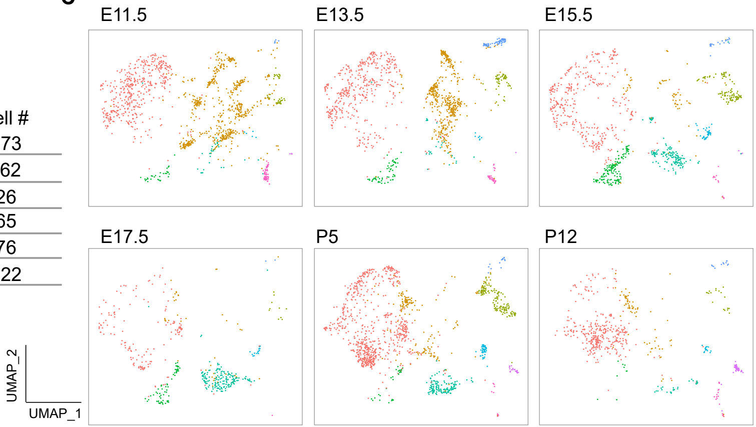

### Supplementary Figure4

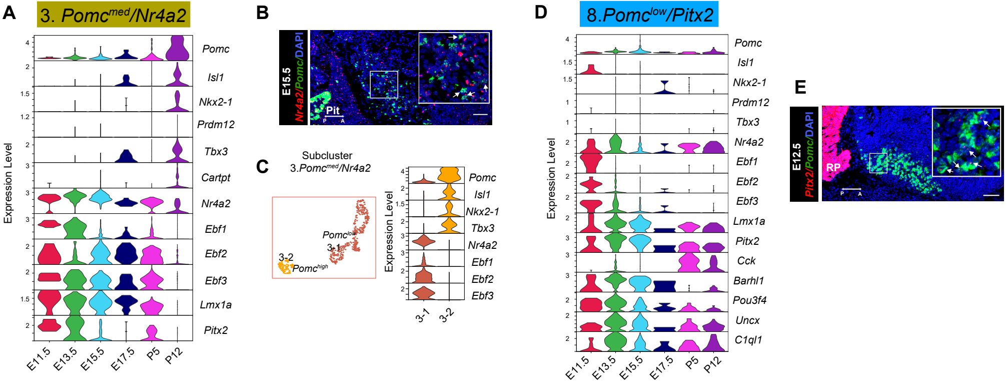

### Supplementary Figure5

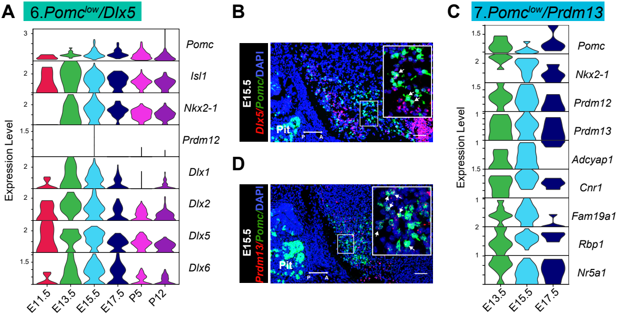

### Supplementary Figure6

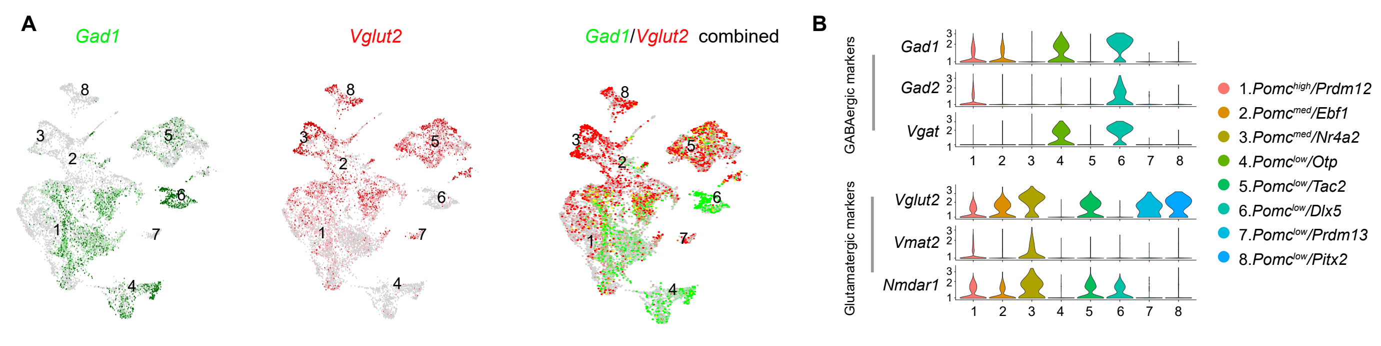

### Supplementary Figure7

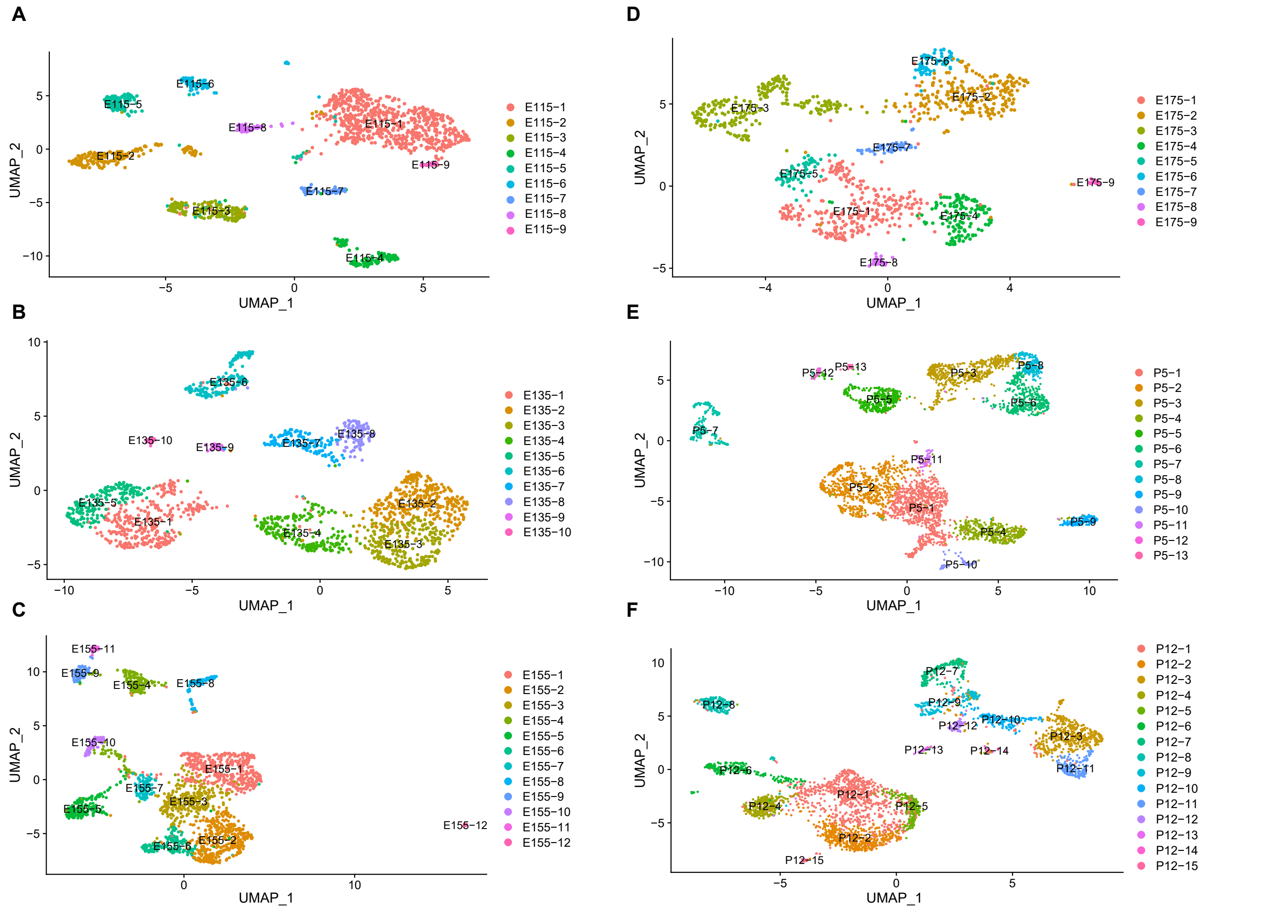

### Supplementary Figure9

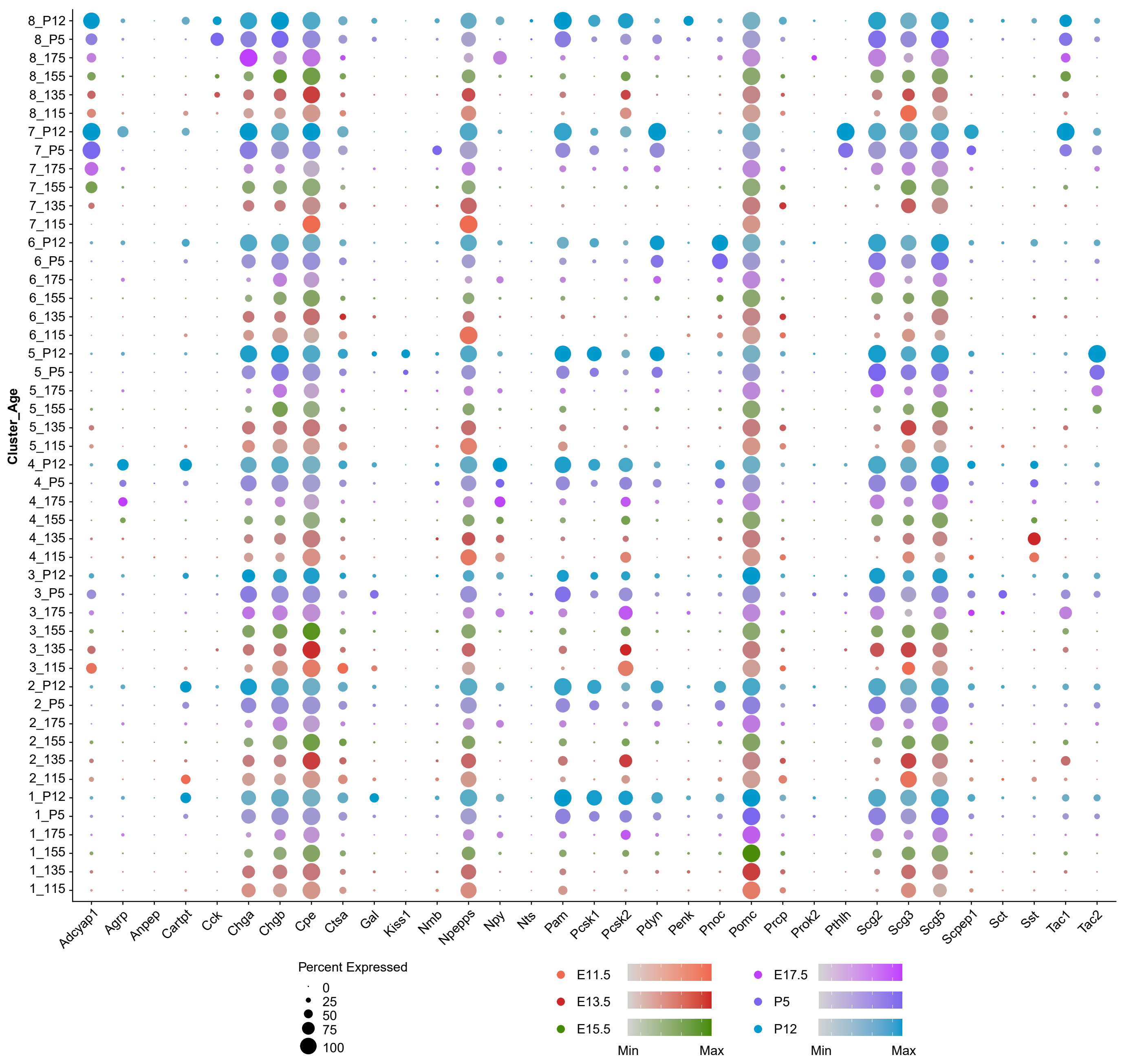

### Supplementary Figure10a

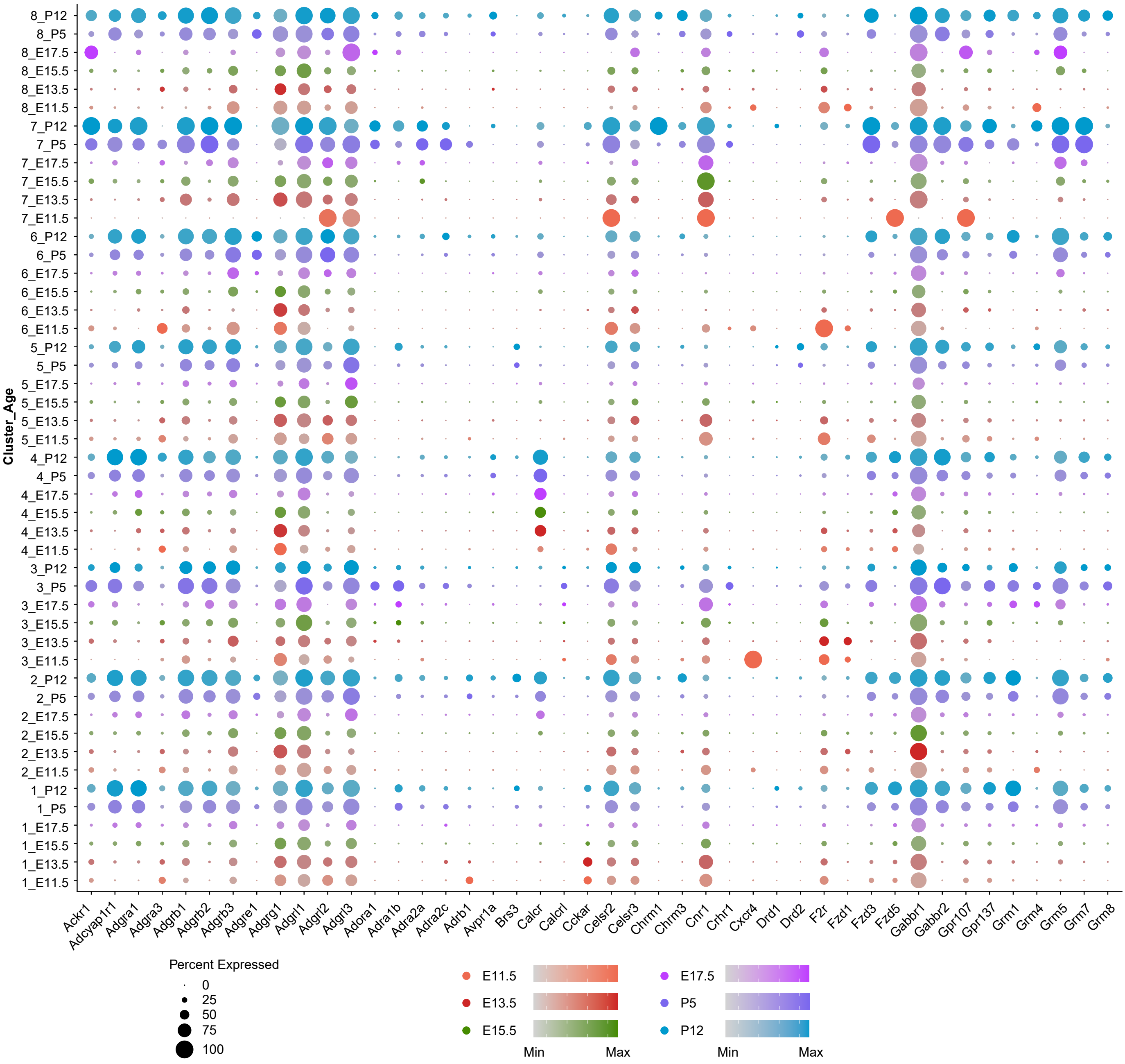

### Supplementary Figure10b

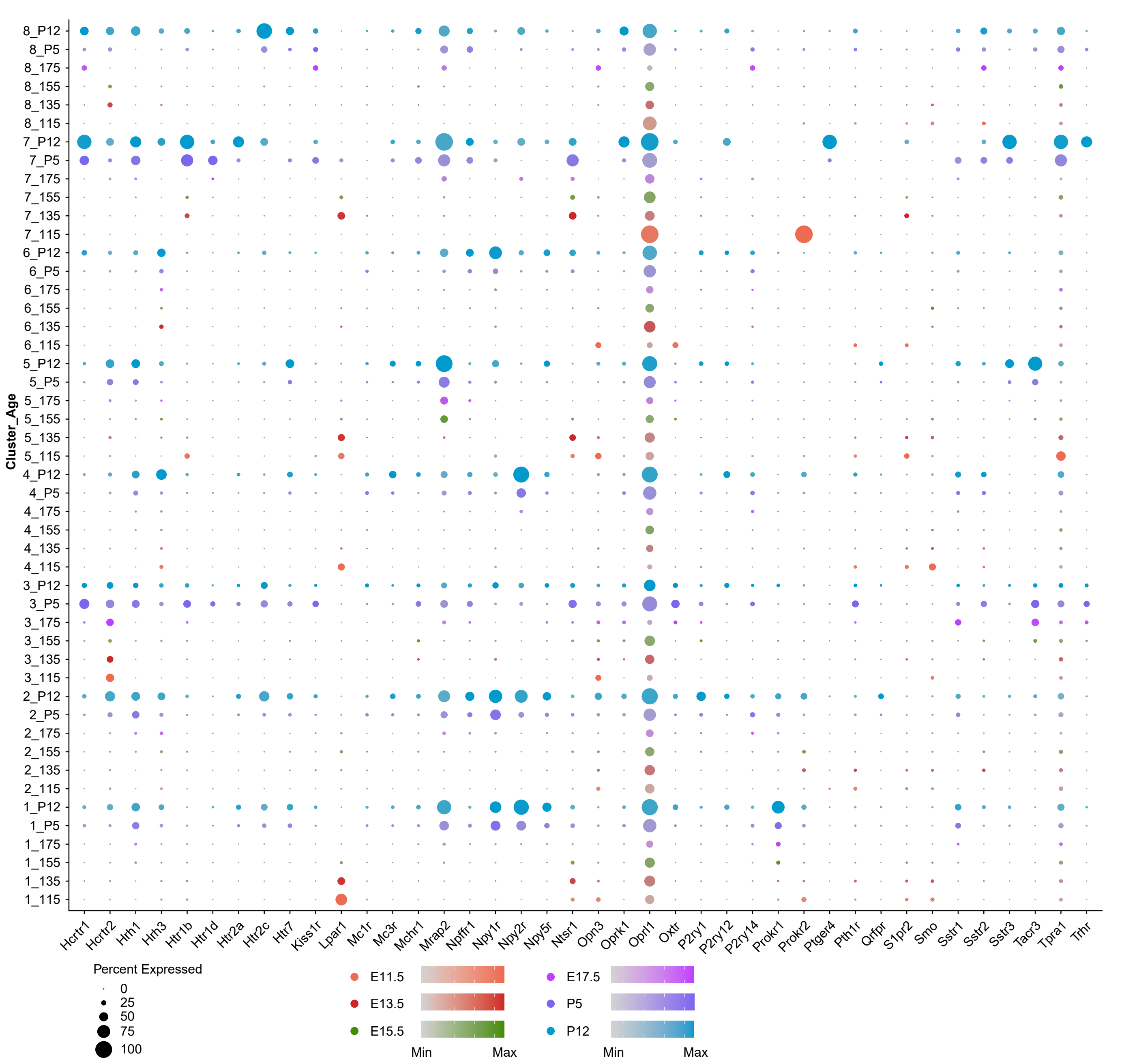

### Supplementary Figure11

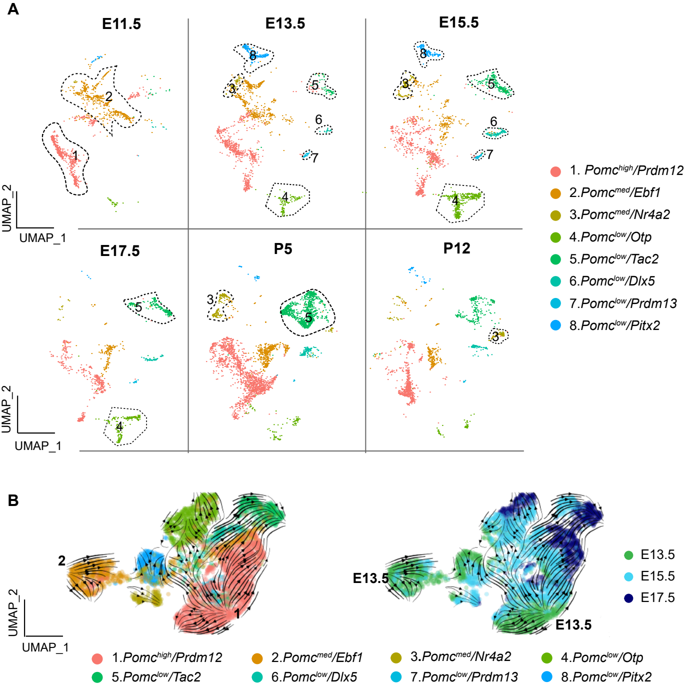

### Supplementary Figure12

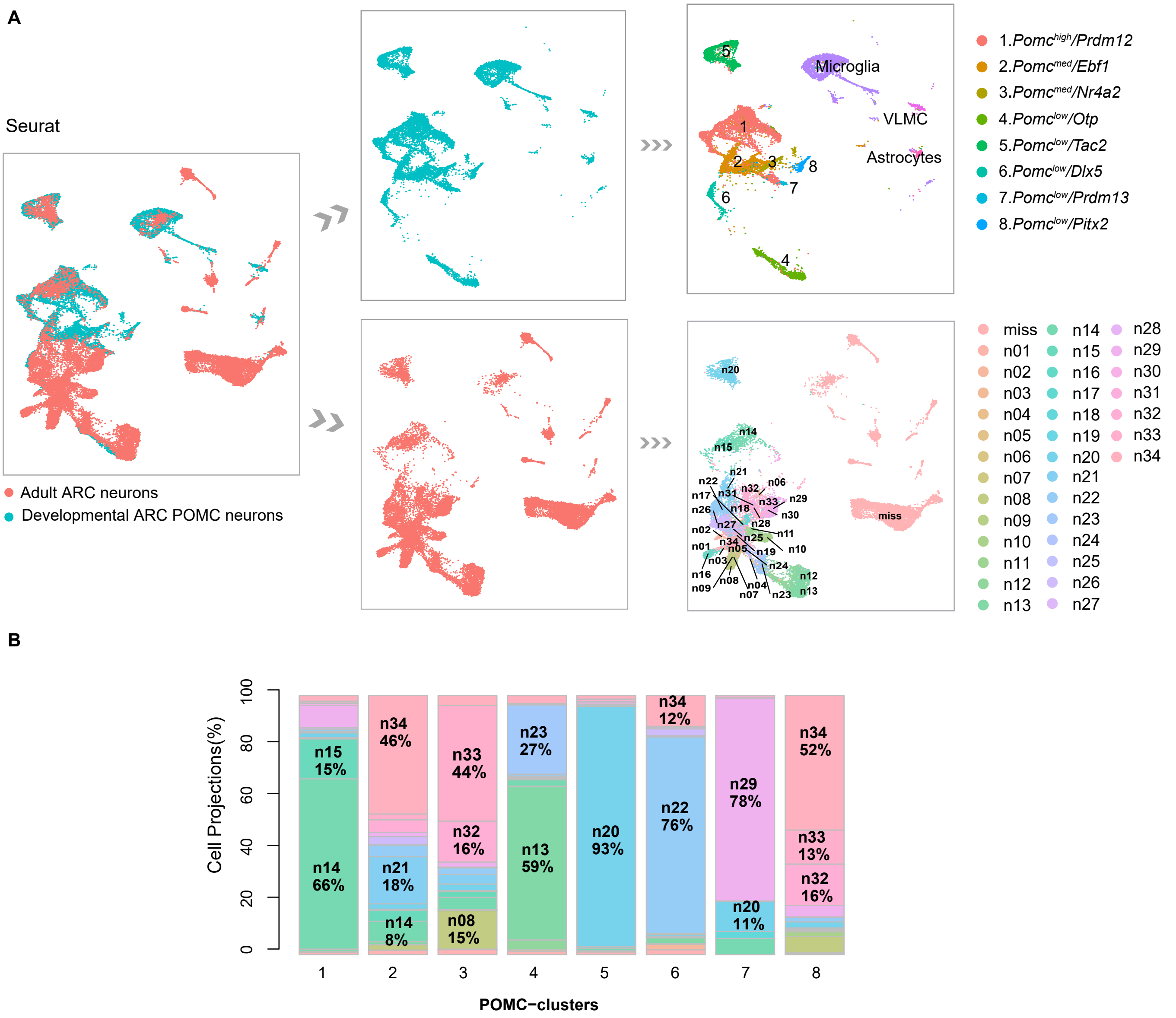

### Supplementary Figure14

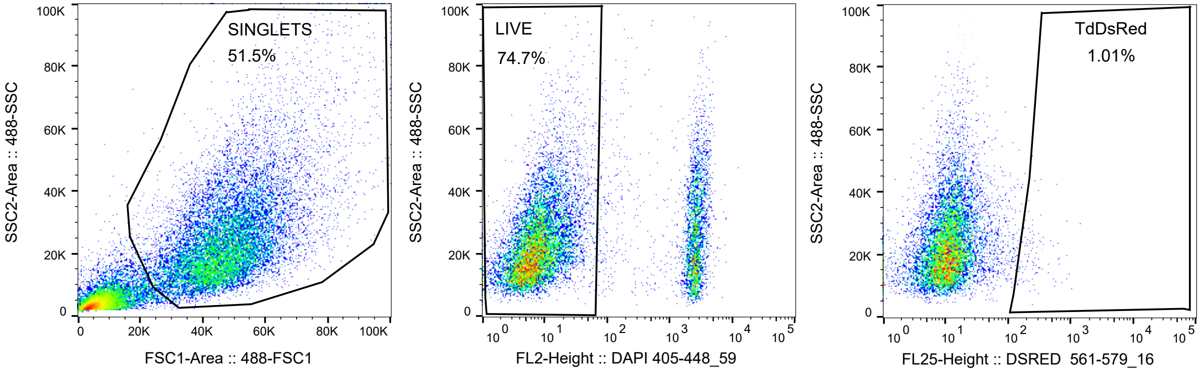
