## Supplementary Figure8 for "Developmental single-cell transcriptomics of hypothalamic POMC progenitors reveal the genetic trajectories of multiple neuropeptidergic phenotypes"

| E11.5 | E13.5 | E15.5 | E17.5 | P5 | P12 | Integrated Clusters |
|---|---|---|---|---|---|---|
| E11.5-1 | E13.5-2<br>E13.5-3 | E15.5-2<br>E15.5-6 | E17.5-4 | P5-1 | P12-3 | *1.Pomc*$^{high}$*/Prdm12* |
| | | E15.5-1 | E17.5-3 | P5-10 | P12-13 | *4.Pomc*$^{low}$*/Otp* |
| E11.5-9 | E13.5-4 | E15.5-3 | E17.5-1 | P5-1<br>P5-2<br>P5-11 | P12-3<br>P12-11<br>P12-14 | *1.Pomc*$^{high}$*/Prdm12* |
| E11.5-5<br>E11.5-6<br>E11.5-2 | E13.5-5<br>E13.5-1 | E15.5-10<br>E15.5-10 | | | | *2.Pomc*$^{med}$*/Ebf1* |
| | E13.5-7<br>E13.5-8 | E15.5-4 | E17.5-9? | P5-7 | P12-9 | *3.Pomc*$^{med}$*/Nr4a2* |
| | E13.5-9 | E15.5-4<br>E15.5-9 | E17.5-9 | P5-7 | P12-9 | *8.Pomc*$^{low}$*/Pitx2* |
| E11.5-4<br>E11.5-8 | E13.5-10 | E15.5-11 | | | | *3.Pomc*$^{med}$*/Nr4a2* |
| | | E15.5-5 | E17.5-2<br>E17.5-6 | P5-8<br>P5-3<br>P5-6 | P12-7<br>P12-7 | *5.Pomc*$^{low}$*/Tac2* |
| | | E15.5-7 | E17.5-7 | P5-9 | P12-12 | *6.Pomc*$^{low}$*/Dlx5* |
| | | | E17.5-5<br>E17.5-8 | P5-4 | P12-10 | *2.Pomc*$^{med}$*/Ebf1* |
| | | E15.5-12 | | P5-5<br>P5-13 | P12-1<br>P12-2<br>P12-5<br>P12-6?<br>P12-8? | 9.Microglia |
| E11.5-3 | E13.5-6 | E15.5-8 | | | P12-6?<br>P12-8? | 10. VLMC |
| | | | | P5-12 | P12-4<br>P12-15 | 11. Astrocytes |
