## Supplementary Figure13 for "Developmental single-cell transcriptomics of hypothalamic POMC progenitors reveal the genetic trajectories of multiple neuropeptidergic phenotypes"

| Developmental POMC$^{+}$ Cells | Adult ARC Neurons |
|---|---|
| ***1.Pomc$^{high}$/Prdm12*** | **n14. *Pomc/Ttr***<br>n15. *Pomc/Anxa2* |
| ***2.Pomc$^{med}$/Ebf1--1*** | **n14. *Pomc/Ttr***<br>**n15. *Pomc/Anxa2*** |
| ***2.Pomc$^{med}$/Ebf1--2*** | **n34. unassigned2**<br>n03. *Th/Sst* |
| ***3.Pomc$^{med}$/Nr4a2--1*** | **n33. unassigned1**<br>n34. unassigned2 |
| ***3.Pomc$^{med}$/Nr4a2--2*** | **n14. *Pomc/Ttr***<br>n15. Pomc/Anxa2 |
| ***4.Pomc$^{low}$/Otp*** | **n13. *Agrp/Gm8773***<br>n23. *Sst/Unc13c* |
| ***5.Pomc$^{low}$/Tac2*** | **n20. *Kiss1/Tac2*** |
| ***6.Pomc$^{low}$/Dlx5*** | **n22. *Tmem215***<br>n34. unassigned2 |
| ***7.Pomc$^{low}$/Prdm13*** | **n29. *Nr5a1/Adcyap1***<br>n20. *Kiss1/Tac2* |
| ***8.Pomc$^{low}$/Pitx2*** | **n34. unassigned2**<br>n32. *Slc17a6/Trhr* |
